# PINK1/Parkin-dependent mitophagy mediates astrocytic inflammatory responses to mitochondrial damage

**DOI:** 10.64898/2026.05.11.724378

**Authors:** Julia F. Riley, Charity V. Robbins, Erika L. F. Holzbaur

## Abstract

Astrocytes directly influence neuronal survival and increasingly are understood to contribute to the progression of neurodegenerative diseases including Parkinson’s disease (PD). Mitochondrial damage is a hallmark of PD pathology in both neurons and astrocytes. Damaged mitochondria are cleared by PINK1/Parkin-mediated mitophagy; loss-of-function mutations in either PINK1 or Parkin are sufficient to cause PD. Neuronal mitophagy is well-studied, but far less is known about how mitochondrial dysfunction in astrocytes affects neural health. While microglial release of pro-inflammatory cytokines has been shown to induce astrocytes to mount their own inflammatory response, we hypothesize that a more direct pathway is involved, and that mitochondrial damage to astrocytes directly triggers release of proinflammatory cytokines. To address these questions, we treated primary murine cortical astrocytes with oxidative phosphorylation (OXPHOS) inhibitors antimycin A (AA) and oligomycin A (OA) and observed the PINK1-dependent accumulation of Parkin on damaged mitochondria, leading to phospho-ubiquitination of proteins in the outer mitochondrial membrane and the recruitment of the autophagy receptor SQSTM1/p62. To identify transcriptional changes caused by mitochondrial damage and the resulting activation of mitophagic machinery, we performed bulk RNA-sequencing on astrocytes isolated from WT, PINK1^-/-^, or Parkin^-/-^ mice treated with AA/OA or a vehicle control. In WT astrocytes, TNF-α signaling via NF-κB was the most significantly upregulated pathway following OXPHOS inhibition. OXPHOS inhibitor treatment also stimulated p62 expression, while NF-κB inhibition prevented this upregulation. Astrocytic secretion of cytokines, including TNF-α, was increased following mitochondrial damage; this secretion was dependent on NF-κB activation and occurred at levels sufficient to induce mitochondrial depolarization in hippocampal neurons. Compared to WT astrocytes, PINK1^-/-^ astrocytes showed a significant reduction in transcriptional signatures associated with TNF-α signaling following mitochondrial damage, while Parkin^-/-^ astrocytes exhibited upregulation of both IFN-γ and IFN-α signaling. These findings indicate altered inflammatory responses to mitochondrial damage in the absence of functional PINK1 or Parkin. Finally, we analyzed scRNA-sequencing data from substantia nigra astrocytes harvested from human brain tissue from PD-positive or control samples. Distinct clusters comprised predominantly of PD-positive or control astrocytes emerged. Astrocytes in the PD-positive cluster were enriched for NF-κB, IFN-α and IFN-γ responses, consistent with the signaling observed *in vitro* post-OXPHOS inhibition. Together, these findings identify inflammatory signatures activated by mitochondrial damage in astrocytes, and establish this pathway as a potential contributor to neuroinflammation in PD.

## INTRODUCTION

Mitochondrial function is critical to cellular energy metabolism, but radical intermediates produced during oxidative phosphorylation (OXPHOS) make these organelles particularly vulnerable to oxidative damage^1,2^. This is especially true in long-lived cells with high metabolic demands, including neurons and some glial cell subtypes such as astrocytes^3–7^. Astrocytic OXPHOS is critical for neuronal health^8^, and mitochondria in astrocytes produce higher levels of reactive oxygen species than neuronal mitochondria^7^. Multiple mechanisms have evolved to maintain mitochondrial homeostasis, including removal of damaged mitochondria via a selective form of autophagy known as mitophagy^9–12^. Mutations that disrupt mitophagy are causal for the neurodegenerative disorders Parkinson’s disease (PD) and amyotrophic lateral sclerosis (ALS), highlighting the importance of this process for maintaining a healthy nervous system^13–18^. While direct mitochondrial damage to neurons is sufficient to induce neuronal death^19–22^, the impact of mitochondrial dysfunction on astrocytes, which are indispensable for maintaining an environment favorable to neuronal survival^23–26^, is less understood.

Astrocytes undergo mitochondrial damage in PD to an extent that is comparable with the damage that occurs in neurons^27–30^ . Previous work has shown that microglia can initiate inflammatory signaling that can then be propagated by astrocytes, leading to neurotoxicity^31^. However, it is much less clear if astrocytes can initiate proinflammatory signaling cascades in response to mitochondrial stress. Furthermore, the consequences that mitochondrial damage and/or defective mitophagy in astrocytes have for neuronal health have not been investigated. Given accumulating evidence that astrocytes exhibit mitochondrial dysfunction during aging that is exacerbated in diseases like PD^27,29,30,32^, identifying astrocytic responses to mitochondrial damage is fundamental to understanding how astrocytes contribute to both normal inflammaging and to neurodegenerative disease progression.

Oxidative damage to mitochondria is well-established to activate PINK1/Parkin-dependent mitophagy in neurons, and has recently been observed in astrocytes as well^33–36^. The coordinated actions of PTEN-Induced Kinase 1 (PINK1) and the E3 ubiquitin ligase Parkin result in the construction of phospho-ubiquitin chains on proteins in the outer mitochondrial membrane (OMM)^12,34^, which in turn leads to the recruitment of ubiquitin-binding autophagy receptors to damaged mitochondria^12^. Autophagy receptors facilitate engulfment of damaged mitochondria into autophagosomes, thus leading to isolation and subsequent degradation of damaged mitochondrial fragments^15,37^. Concurrent with the recruitment of autophagy receptors to damaged mitochondria, our group has demonstrated that the PINK1/Parkin-dependent ubiquitination of OMM proteins is also sufficient to directly activate the Nuclear Factor-κB (NF-κB) innate immunity pathway^38^. The NF-κB effector molecule (NEMO) is rapidly recruited to damaged mitochondria in multiple cell types^38,39^, including astrocytes^38^, and acts as a scaffold for the IκB kinase (IKK) signaling complex that initiates NF-κB signaling. These findings raise the possibility that initiation of mitophagy in astrocytes directly activates inflammatory cascades that may detrimentally affect neuronal health.

Here, we examined the consequences of OHPHOS failure on murine cortical astrocytes, which we show is characterized by the recruitment of the autophagy receptor SQSTM1/p62. Bulk RNA sequencing analysis validated by western blotting indicates that NF-κB signaling is a prominent aspect of the WT astrocyte response to oxidative damage. This response leads to a significant upregulation of TNF-α transcription, which is blocked by treatment of astrocytes with a small molecule inhibitor of NF-κB. Cytokine profiling demonstrates that oxidative damage to astrocytes induces an NF-κB-dependent secretory response that includes TNF-α release. Treatment of hippocampal neurons with corresponding levels of TNF-α was sufficient to negatively impact mitochondrial polarization and to alter mitochondrial network morphology, indicating that oxidative stress to astrocytes is sufficient to promote neuronal dysfunction. NF-κB activation also induces upregulation of the autophagy adaptor SQSTM1/p62, facilitating efficient mitochondrial clearance. Importantly, this NF-κB-dependent upregulation constitutes a negative feedback loop that temporally limits damage-induced NF-κB signaling. In contrast to these findings from wild type astrocytes, transcriptomic analysis of astrocytes from PINK1^-/-^ mice indicates that the damage-induced NF-κB signature is blunted, while astrocytes lacking Parkin exhibit activation of alternative inflammatory signaling pathways, most notably IFN-α and IFN-γ responses. To explore the relevance of these findings to human disease, we analyzed single-cell sequencing data from substantia nigra astrocytes derived from PD-affected human tissue. Strikingly, we identified indicators of IFN-α and IFN-γ signaling parallel to the transcriptional signatures observed in Parkin^-/-^ astrocytes subjected to oxidative damage. These findings demonstrate that oxidative damage in astrocytes elicits a self-limiting inflammatory response modulated by mitophagy; key aspects of the transcriptional signature we observed *in vitro* can be seen in transcriptomic analysis of astrocytes from patients with PD. Together, these findings establish a novel link between the mitochondrial damage response in astrocytes and the inflammatory signatures seen in PD pathology.

## RESULTS

### Mitochondrial damage to astrocytes activates NF-κB-dependent upregulation of TNF-α

Mitochondrial damage to neurons has been implicated in the onset of neurodegeneration in Parkinson’s disease, but the effects of mitochondrial damage to astrocytes, and the contributions of this damage to neuronal loss, are much less understood. While microglial release of pro-inflammatory cytokines has been shown to induce astrocytes to mount their own inflammatory response, we hypothesize that there is likely to be a more direct pathway involved– that mitochondrial damage to astrocytes can directly trigger release of proinflammatory cytokines. To test this hypothesis, we first established that mitochondrial damage elicits a robust PINK1/Parkin-dependent response in astrocytes (Extended Data Fig. 1A). Mitochondrial damage stimulates phospho-ubiquitination of mitochondria treated with the OXPHOS inhibitors Antimycin A (AA) and Oligomycin A (OA) in primary mouse astrocytes, similar to prior observations in rat astrocytes^34^. We observed progressive accumulation of S65 phospho-ubiquitin on damaged mitochondria in wild-type (WT) murine cortical astrocytes via western blotting (Extended Data Fig. 1B, C) and immunocytochemistry (Extended Data Fig. 1D-F). Consistent with the established pathway for PINK1/Parkin-dependent mitophagy, YFP-Parkin is recruited to damaged mitochondria in both monocultured astrocytes (Extended Data Fig. 1E, G) and astrocytes co-cultured with human iPSC-derived neurons (Extended Data Fig. 1H). In contrast, in astrocytes isolated from PINK1^-/-^ mice, we observed mitochondrial fragmentation but saw no YFP-Parkin recruitment following AA/OA treatment (Extended Data Fig. 1I-K).

To assay the effects of mitochondrial damage on astrocytes, we performed bulk RNA sequencing on primary mouse astrocytes treated with either AA and OA or vehicle control for eight hours (Fig. 1A). The most significantly upregulated KEGG pathway observed in response to mitochondrial damage was TNF-α signaling (Fig. 1B). We also performed Gene Set Enrichment Analyses (GSEA) for each of the 50 hallmark gene sets available in the Molecular Signatures Database (MSigDB). The most enriched pathway was TNF-α signaling via NF-κB (Fig. 1C), corroborating these findings and suggesting NF-κB activation as the driving force behind TNF-α production. In addition to TNF-α signaling, we noted enrichment of other inflammatory pathways, including IL-17 signaling, cytosolic DNA-sensing pathway (potentially indicative of cGAS/STING involvement), and pathways associated with viral immune responses, including viral protein interaction with cytokine and chemokine receptor and Influenza A (Fig. 1B). We also saw downregulation of fatty acid metabolism (Fig. 1C), consistent with observations suggesting that mitochondrial damage to astrocytes disrupts their ability to metabolize fatty acids in the brain^44^.

**Figure 1.**
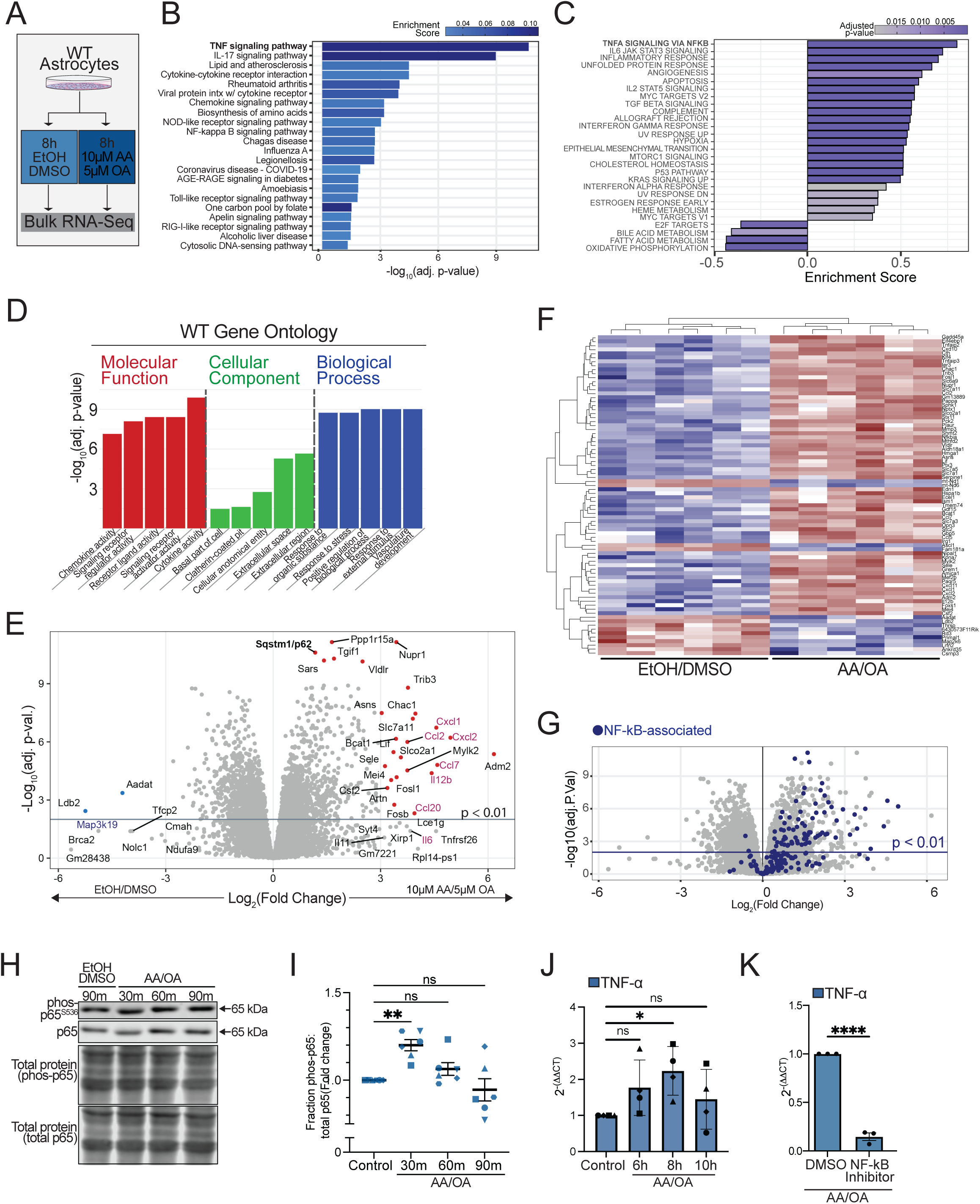
Mitochondrial damage to astrocytes activates NF-κB-dependent upregulation of TNF-α. A) Schematic for bulk-RNA sequencing experiment. B) KEGG terms from gene ontology (GO) analysis comparing transcriptomes of WT astrocytes following 8h treatment with 10 μM AA/ 5 μM OA to those of WT astrocytes treated with a vehicle control (n=6). C) GSEA analysis of all hallmark gene sets significantly different in WT astrocytes following 8 hours 10 μM AA/ 5 μM OA treatment relative to the 8h EtOH/DMSO vehicle control arranged in order of descending enrichment score. D) Top five most significant Molecular Function, Cellular Component, and Biological Processes GO terms enriched in WT astrocytes post-AA/OA compared to vehicle control (n=6). E) Volcano plot of differentially expressed genes (DEGs) in WT astrocytes following 10 μM AA/ 5 μM OA AA/OA treatment (positive log_2_FC) compared to the vehicle control (negative log_2_FC). Transcripts with |log2(FC)| > 3 or a log10(adj. p. value) > 9 are labeled. Dots corresponding to transcripts with an adj. p-value <0.01 and a log2(FC) > 3 or a log10(adj. p-value) >9 are red; dots corresponding to transcripts with an adj. p-value < 0.01 and a log2(FC) < -3 are blue. Selected chemokines and cytokines are shown in pink (n=6). F) Heatmap of differentially expressed genes (DEGs) comparing WT astrocytes following 8h 10 μM AA/ 5 μM OA compared to the vehicle control. Columns represent biological replicates (AA/OA, n=6; control, n=6). Expression values were scaled by row, and adjacent phylograms represent hierarchal clustering of genes (left) and samples (top). G) Volcano plot in (E) of DEGs in WT astrocytes following AA/OA treatment (positive log_2_FC) compared to vehicle control (negative log_2_FC); genes included in TNF-α Signaling via NF-κB pathway in the Hallmark gene collection on the MSigDB are highlighted in blue (n=6). H, I) Quantification of phospho-p65: total p65 ratio in astrocytes treated with 10 μM AA/ 5 μM OA for 30, 60, or 90 minutes or for 90 minutes with vehicle. Each is normalized to the vehicle control for the respective biological replicate (n= 6, ordinary one-way ANOVA with multiple comparisons, error bars indicate mean +/- SEM). J) qPCR measuring transcript levels for *Tnfa* normalized to *Actb* following 6h, 8h, or 10h treatment with 10 μM AA/ 5 μM OA or 10h treatment with vehicle (n= 4, non-parametric ANOVA, error bars indicate mean +/- SEM). K) qPCR measuring transcript levels for *Tnfa* normalized to *Actb* following 8h treatment with 10 μM AA/ 5 μM OA in astrocytes pre-treated with either the IKK-β inhibitor BMS345541 or vehicle control for 48 hours (n=3, unpaired t-test, error = mean +/- SEM). Biological replicates are defined as independent experiments performed on astrocytes from different dissections. For all graphs, p < 0.05 = *; p < 0.01 = **; p < 0.001 = ***; p < 0.0001 = ****.

To further characterize pathways activated following mitochondrial damage in astrocytes, we performed a gene ontology analysis. Molecular function analysis revealed that many of the upregulated transcripts encode cytokines or chemokines (Fig. 1D). Transcripts associated with cellular stress responses and the extracellular space were also upregulated, further implicating inflammatory signaling. Upon direct analysis of specific DEGs upregulated following OXPHOS inhibition in astrocytes, we found that many of these transcripts encode signaling molecules affiliated with inflammatory signaling. This included several C-X-C motif chemokines (e.g. Cxcl1, Cxcl2 and Cxcl10) and C-C motif chemokines (e.g. Ccl20) (Fig. 1E, F).

As NF-κB and TNF-α signaling-associated terms were among the most significantly affected pathways in our transcriptomic analysis, we asked how these changes might influence inflammatory signaling in astrocytes. Genes annotated as being involved in TNF-α signaling via NF-κB were enriched post-AA/OA treatment compared to the vehicle control (Fig. 1F, G). Specifically, NF-κB-associated genes such as TNF-α-induced protein 2 (TNFAIP2) and matrix metalloprotease 3 (MMP3)^40^ were among the DEGs consistently upregulated post-stress (Fig. 1F). Collectively, these data suggest a robust role for NF-κB signaling in initiating inflammatory signaling stimulated by mitochondrial damage to astrocytes. To further validate activation of NF-κB following mitochondrial damage, we performed western blotting experiments to detect phosphorylation of the major NF-κB transcription factor, p65 (also called RelA). Astrocytes were treated for 30, 60, or 90 minutes with AA/OA or vehicle control. We detected a significant increase in the phosphorylated fraction of p65 (Fig. 1H, I) within 30 minutes of AA/OA treatment, indicating rapid and transient activation of NF-κB on a timescale consistent with direct activation of the pathway. We next performed qPCR and found that TNF-α transcription peaks 8 hours post-AA/OA treatment (Fig.1J). Transcription of TNF-α then begins to decline, suggesting a possible negative feedback loop to prevent chronic elevation of NF-κB signaling following mitochondrial damage. To confirm that AA/OA-induced TNF-α production is NF-κB-mediated, we treated astrocytes with a small molecule inhibitor of IKKβ, a component of the tripartite protein complex that canonically initiates NF-κB-mediated signaling. We observed an 85% decrease in TNF-α transcript levels post-AA/OA treatment when IKKβ was inhibited with BMS345541, confirming that TNF-α production in response to mitochondrial damage is largely mediated by NF-κB signaling (Fig. 1K).

### NF-κB-mediated TNF-α secretion by astrocytes induces mitochondrial depolarization and loss of mitochondrial volume in neurons

Given the major role of NF-κB activation in mediating TNF-α transcription following mitochondrial damage to astrocytes (Fig. 1K), we asked whether mitochondrial damage activates the astrocytic secretion of cytokines and chemokines in an NF-κB-dependent manner. We treated astrocytes with AA/OA, AA/OA in combination with BMS345541, or vehicle control and performed a cytokine screen on astrocyte-conditioned media to detect the presence of 44 different cytokines, chemokines, and growth factors (Fig. 2A, Table S1). We found that there was a striking reduction in TNF-α secretion when NF-B signaling was inhibited. The secretion of many other cytokines and chemokines was downregulated by NF-κB inhibition, including CX3CL1/Fractalkine, known for its role in recruiting microglia to the site of injury. This indicates that NF-κB is a central mediator of astrocyte secretory responses following mitochondrial damage. While the production of several cytokines increased following AA/OA treatment, the secretion of interferon gamma (IFNγ) was reduced following AA/OA compared to vehicle control (Figure 2A). Together, these data establish NF-κB as a major mediator of astrocytic signaling in response to cell-intrinsic stress.

**Figure 2.**
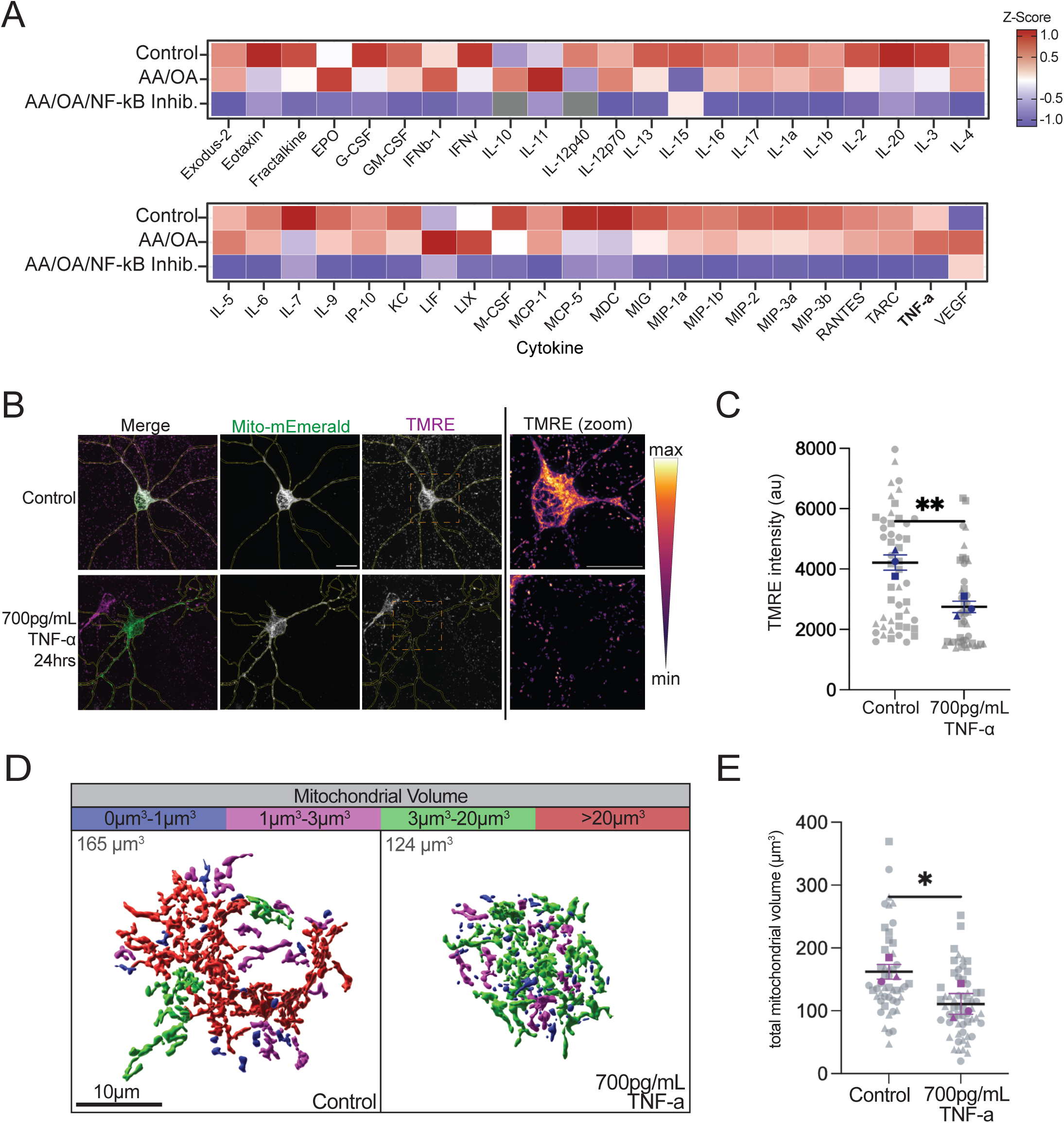
NF-κB-mediated TNF-α secretion by astrocytes induces mitochondrial depolarization and loss of mitochondrial volume in neurons. A) Cytokine panel performed on conditioned media from WT astrocytes treated for 48 hours with the IKK-β inhibitor BMS345541 and then an additional 16 hours with 10 μM AA/OA or a vehicle control. Normalized z-scores indicate relative levels of secreted factors calculated for each peptide based on the average across six independent biological replicates (n=6). Gray squares indicate conditions in which all six biological replicates fell below the detectable range. B) TMRE staining in rat hippocampal neurons (DIV14) transfected with mitochondrially-targeted mEmerald and treated with 700 pg TNF-α or vehicle control for 24 hours. Representative max projections are shown. Scale bars = 20 μm. C) Mitochondrial TMRE intensity quantification. Gray dots represent individual cell measurements; blue dots represent average intensity for independent biological replicates (n= 3, unpaired t-test, error = mean +/- SEM). D) Volume renderings of HSP60 staining in the soma of rat hippocampal neurons (DIV14) treated with 700 pg/mL TNF-α or vehicle control for 24 hours. Mitochondrial objects are colorized based on their volume. Scale = 10 μm. E) Total mitochondrial volume in hippocampal rat neuron somas following 24h TNF-α treatment. Gray dots indicate individual soma measurements, purple dots indicate average volume for each independent biological replicate (n=3, paired t-test, error = mean +/- SEM). Biological replicates are defined as independent experiments performed on astrocytes or neurons from different dissections. For all graphs, p < 0.05 = *; p < 0.01 = **.

Profiling of conditioned media from astrocytes treated with AA and OA indicated that astrocytes secreted ∼700pg/mL TNF-α following 8h mitochondrial damage (Table S1). We therefore asked whether this concentration of TNF-α could impact mitochondrial health in neurons. We treated rat hippocampal neurons with 700 pg/mL TNF-α for 24h and assessed mitochondrial membrane potential using tetramethyl rhodamine (TMRE). Mitochondrial TMRE intensity was significantly decreased in neurons following 24h TNF-α treatment compared to a vehicle control (Fig. 2B, C). In parallel, we analyzed changes in mitochondrial morphology in neurons following treatment with TNF-α. After a 24h treatment with either 700pg/mL TNF-α or vehicle control, we performed immunocytochemistry to map the somatic mitochondrial network in 3D. The overall volume of mitochondria in the soma was significantly decreased following TNF-α treatment (Fig. 2D, E), suggesting loss of mitochondrial content, potentially due to enhanced turnover. These results indicate that exposure to TNF-α at a concentration akin to what is released by astrocytes that have undergone OXPHOS failure is sufficient to directly influence the health of the mitochondrial network in neurons.

### NF-κB signaling mediates p62 upregulation and mitochondrial clearance upon astrocytic mitochondrial damage

The transient nature of the upregulation of TNF-α transcription induced by OXPHOS inhibition in astrocytes suggested to us that there might be a negative feedback loop preventing prolonged inflammatory signaling. Previous work from our group has shown that one mechanism leading to NF-κB activation in response to mitochondrial damage involves recruitment of the NF-κB effector molecule (NEMO) to ubiquitinated proteins in the outer mitochondrial membrane produced by PINK1-dependent activation of the E3 ligase Parkin^38^. Specifically, activation of PINK1/Parkin-dependent mitophagy in astrocytes was sufficient to recruit NEMO to ubiquitinated mitochondria^38^. Efficient induction of mitophagy would be predicted to induce degradation of damaged mitochondria, leading to attenuation of NF-κB signaling generated via this mechanism and acting as a negative feedback loop to prevent chronic inflammatory signaling.

Interestingly, the ubiquitin-binding autophagy receptor Sqstm1/p62 (hereafter referred to as p62) was among the most significantly upregulated genes following AA/OA treatment in WT astrocytes, upregulated ∼2.3-fold relative to vehicle control (Fig. 3A). The only other autophagy receptor that was significantly upregulated following AA/OA treatment was OPTN, whose expression relative to baseline increased ∼1.5-fold following AA/OA compared to vehicle control (Extended Data Fig. 2A). We confirmed increased levels of p62 following 24h AA/OA treatment in astrocytes via western blot (Fig. 3B, C). We also established that this treatment is sufficient to elicit PINK1-dependent clearance of damaged mitochondria, as is evidenced by the degradation of the mitochondrial membrane protein Mic60 and other mitophagy-associated proteins such as Parkin and Mitofusin 2 (Fig. 3B, C). While p62 is not required for mitophagy in HeLa cells^12,15^, we wondered if p62 might play a more specific role in clearing damaged mitochondria in astrocytes. To test this possibility, we again treated astrocytes with AA/OA or vehicle while concurrently inhibiting autophagic degradation using the small molecule inhibitor Bafilomycin A (BafA). We then quantified levels of known autophagy receptors. These include Optineurin (OPTN), Tax1-binding protein 1 (TAX1BP1), and p62. Consistent with effective activation of mitophagic flux, we observed significant accumulation of the autophagy marker LC3II in BafA-treated cells (Fig. 3D). Of the receptors examined, p62 exhibited the greatest accumulation following AA/OA/BafA treatment compared to AA/OA alone (Fig. 3D, Extended Data Fig. 2B, C). Using ICC, we found that endogenous p62, known to form biomolecular condensates, forms pronounced puncta following AA/OA treatment in astrocytes (Fig. 3E, F). On average, 51% of these puncta colocalized with phospho-ubiquitin. To determine whether p62 interacts with phospho-ubiquitin on the mitochondrial surface, we performed a proximity ligation assay (PLA) to detect proximity of p62 (<40nm) to phospho-ubiquitin, indicative of protein-protein interactions. We detected a significant increase in the proximity between the two proteins following AA/OA-induced mitochondrial damage compared to treatment with the vehicle control (Fig. 3G, H). Overall, these data suggest that p62 facilitates mitophagic degradation of damaged mitochondria in astrocytes (Extended Data Fig. 2D).

**Figure 3.**
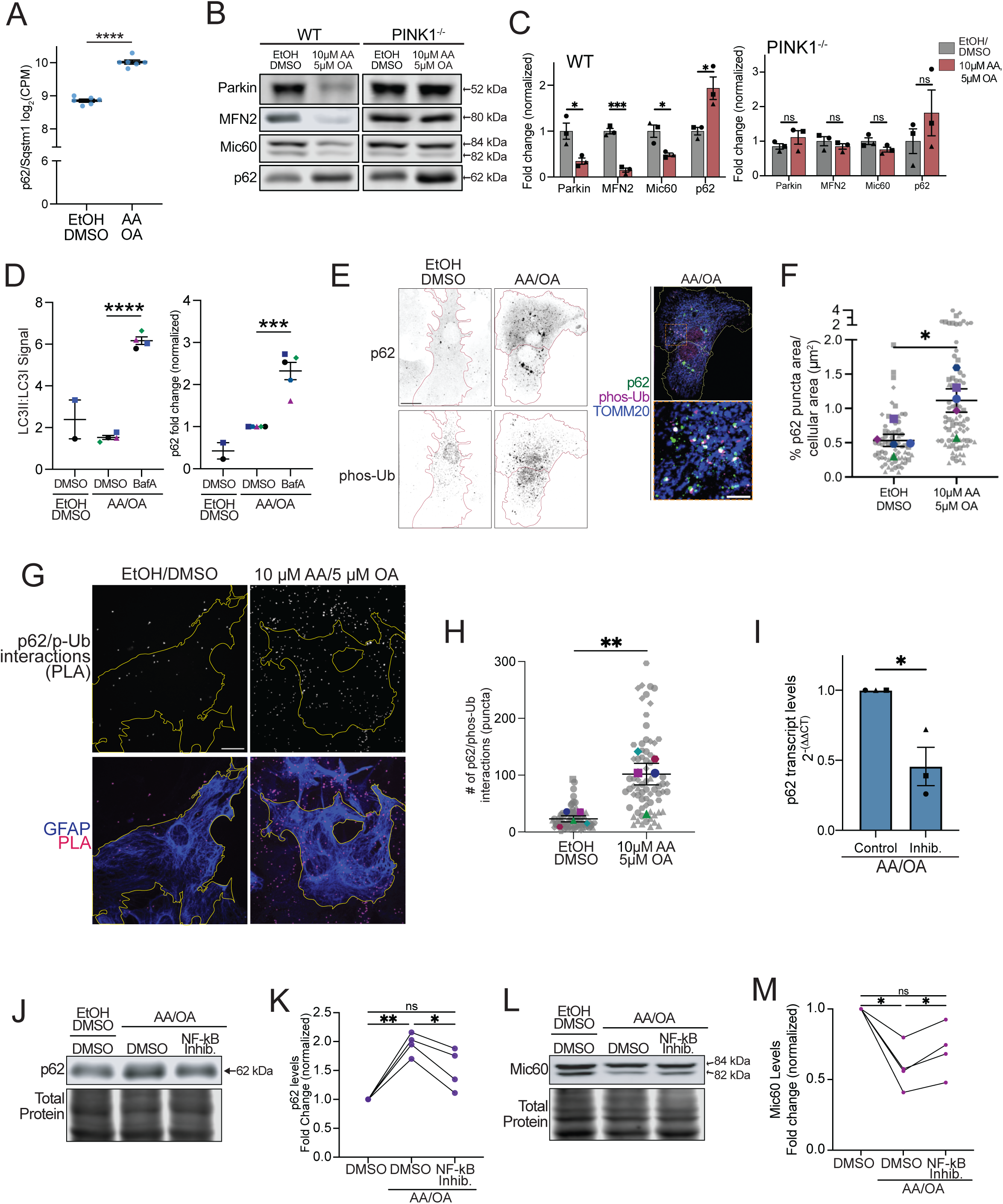
NF-κB signaling mediates p62 upregulation and mitochondrial clearance upon astrocytic mitochondrial damage. A) Log_2_CPM for p62 in WT astrocytes post-stress compared to the control (n=6; unpaired t-test; error bars indicate mean +/- SEM). B) Representative blots for WT and PINK1 KO astrocytes treated with 10 μM AA and 5 μM OA or vehicle control for 24h. C) Western blots for Parkin, MFN2, Mic60, and p62 levels in WT astrocytes (left) or PINK1^-/-^ astrocytes (right) following 24h 10 μM AA/ 5 μM OA compared to cells treated with vehicle control. Protein levels normalized to average for respective controls (n=3, unpaired t-tests between vehicle control and AA/OA-treated condition for each protein of interest, error = mean +/- SEM). D) Western blots for LC3II:LC3I ratio (left) and p62 levels (right) in WT astrocytes treated for 16h with either 10nM Bafilomycin A (BafA) or vehicle control in conjunction with either 10 μM AA/ 5 μM OA or vehicle control. Each replicate is normalized to the AA/OA-only condition (n=2 for treatment with both vehicle controls, n=5 for both conditions with AA/OA; one-way ANOVA with multiple comparisons, error= mean +/- SEM). E,F) p62 and phospho-ubiquitin (S65) immunostaining in WT astrocytes treated with 10 μM AA/5 μM OA or vehicle control for 2 hours. Ectopically expressed mitochondrial TOMM20-Halo is also shown. p62 puncta formation quantified as percent of cell area (n=5, paired t-test, error bars are mean +/- SEM). Representative max projections shown. Scale bar =20 μm. G, H) Proximity ligation assay for p62 and phospho-ubiquitin in astrocytes treated with 10 μM AA/5 μM OA or vehicle control for 2 hours. Representative images (G) and quantification (H) shown. Gray dots represent individual cell measurements; colored dots represent average for each biological replicate. (n=5, unpaired t-test, error bars are mean +/- SEM). Scale bar =20 μm; zoom scale bar = 5 μm. I) qPCR measuring transcript levels for *Sqstm1* (p62) in WT astrocytes normalized to *Actb* following 8h treatment with 10 μM AA/ 5 μM OA in astrocytes pre-treated with either the IKK-β inhibitor BMS345541 or vehicle control for 48 hours (n=3, unpaired t-test, error = mean +/- SEM). J-M) Western blots for p62 (J, K) or Mic60 (L, M) expression in astrocytes treated for 72 hours with the IKK-β inhibitor BMS345541 or a vehicle control, with 10 μM AA/ 5 μM OA or a vehicle control added for the final 24 hours of treatment (n= 4, paired t-test, error = mean +/- SEM).

Given that our DEG analysis had identified p62 as being one of the transcripts most significantly upregulated by mitochondrial damage to WT astrocytes and that NF-κB is robustly activated by mitochondrial damage in these cells, we next sought to determine whether p62 upregulation following mitochondrial damage is facilitated by NF-κB signaling. We treated astrocytes with BMS345541 and either AA/OA or a vehicle control. We found that inhibition of NF-κB led to a significant decrease in p62 transcription following 8h AA/OA treatment in astrocytes (Fig. 3I), as well as a decrease in levels of p62 protein following 24h AA/OA treatment in these cells (Fig. 3J, K). Thus, we asked whether NF-κB inhibition prevented efficient clearance of damaged mitochondria in these cells, and found that treatment with BMS345541 prevented significant clearance of the mitochondrial marker Mic60 following 24h AA/OA treatment in astrocytes compared to treatment with AA/OA alone (Fig. 3L, M). Together, these data indicate that NF-κB activation facilitates efficient clearance of damaged mitochondria in astrocytes, acting as a negative feedback loop to prevent chronic inflammation.

### OXPHOS inhibition triggers p38/MAPK activation and CREB phosphorylation in WT astrocytes

To better understand the nature of the observed inflammatory signatures induced by mitochondrial damage in astrocytes, we performed a GSEA to identify potential transcription factors whose activity is enriched in astrocytes following mitochondrial damage compared to vehicle control. About 75% of the top 50 signatures enriched in astrocytes following mitochondrial damage can be associated with one of five transcriptional motifs: NF-κB, STAT3, STAT5, Activating Transcription Factor/ cAMP-responsive element-binding protein (ATF/CREB, respectively), and AP1 (Fig. 4A, Extended Data Fig. 3A). This suggests that the mitochondrial damage response in astrocytes is nuanced and likely consists of both pro-inflammatory and pro-survival components.

**Figure 4.**
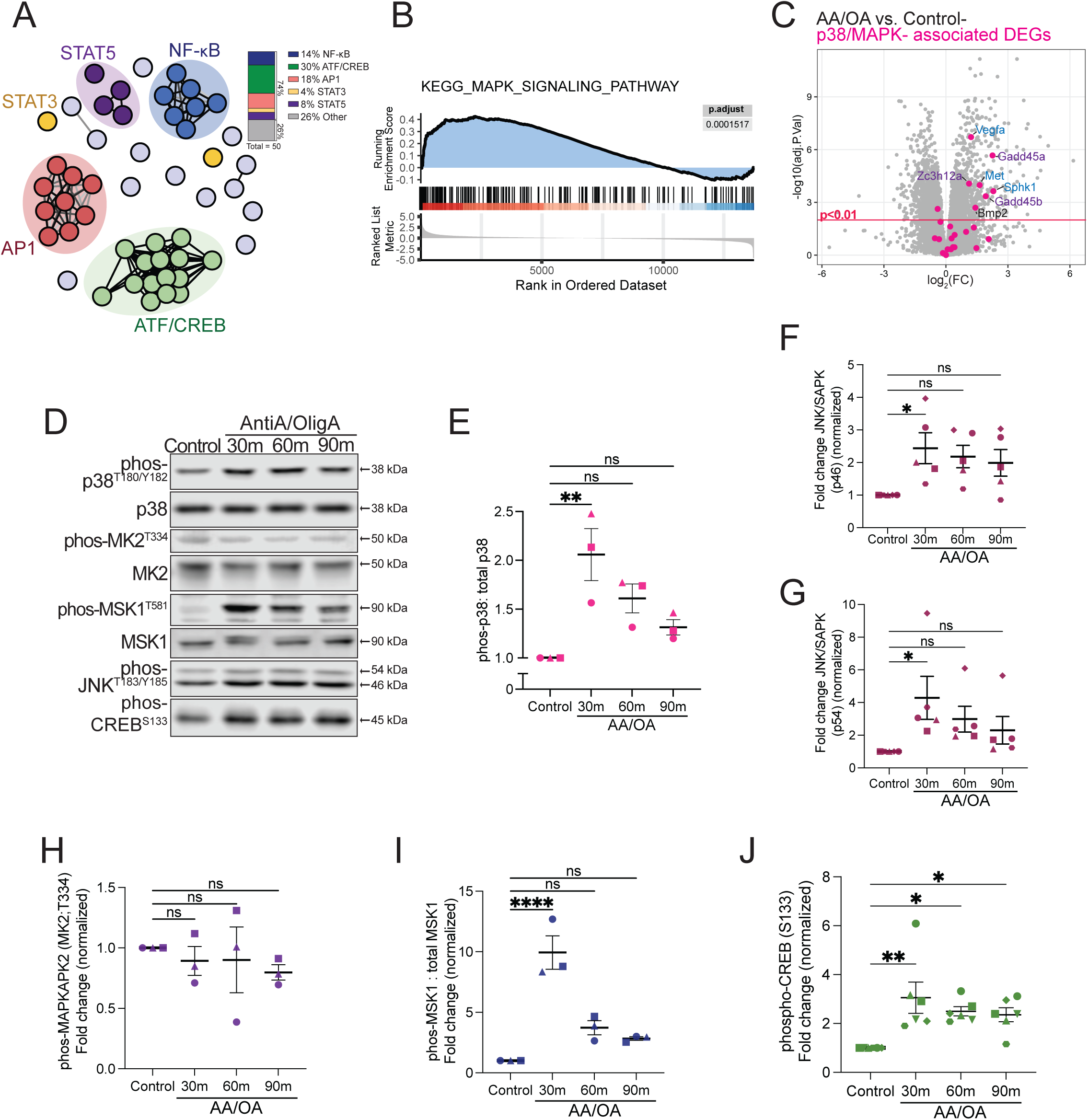
OXPHOS inhibition triggers p38/MAPK activation and CREB phosphorylation in WT astrocytes. A) Top 50 most significantly enriched TFT-Legacy terms in WT astrocytes following AA/OA treatment compared to the control. To simplify results and identify key pathways, terms are arranged according to the percent similarity in genes that define each term. Each circle represents a TFT-legacy term (n=6). B) GSEA for genes listed in the KEGG MAPK signaling pathway shows enrichment of this pathway in WT astrocytes treated with 8h AA/OA compared to those treated with vehicle control (n=6). C) Volcano plot of differentially expressed genes (DEGs) in WT astrocytes following AA/OA treatment (positive log_2_ (Fold Change)) compared to the vehicle control (negative log_2_FC), with dots corresponding with p38/MAPK signaling-associated genes highlighted in magenta. D) Representative western blots for astrocytes treated with 30, 60, or 90 minutes of 10μM AA/ 5μM OA or for 90 minutes with vehicle control. E) Ratio of phospho-p38 (T180/Y182) to total p38 following 30, 60, or 90 minutes of 10μM AA/ 5μM OA or 90 minutes of vehicle control (n= 3, ordinary one-way ANOVA with multiple comparisons, error bars represent mean +/- SEM). F, G) Phospho-JNK/SAPK levels (both the p46 and p54 isoforms; Thr183/Tyr185) following 30, 60, or 90 minutes of 10μM AA/ 5μM OA or 90 minutes of vehicle control (n= 5, ordinary one-way ANOVA with multiple comparisons, error bars represent mean +/- SEM). H) Phospho-MAPKAPK2 (T334) remained undetectable following 30, 60, or 90 minutes of 10μM AA/ 5μM OA or 90 minutes of vehicle control (n= 3; ordinary one-way ANOVA with multiple comparisons; error bars represent mean +/- SEM). I) The ratio of phosphorylated MSK1 (T180/Y182) to total p38 following 30, 60, or 90 minutes of 10μM AA/ 5μM OA or 90 minutes of vehicle control (n= 8, ordinary one-way ANOVA with multiple comparisons, error bars represent mean +/- SEM). J) Phosphorylated CREB levels (S133) following 30, 60, or 90 minutes of 10μM AA/ 5μM OA or 90 minutes of vehicle control (n= 6, ordinary one-way ANOVA with multiple comparisons, error bars represent mean +/- SEM). Biological replicates are defined as independent experiments performed on astrocytes from different dissections. For all graphs, p < 0.05 = *; p < 0.01 = **; p < 0.001 = ***; p < 0.0001 = ****.

One of the transcription factors whose activity was enriched in astrocytes following AA/OA treatment was STAT3 (Fig. 4A, Extended Data Fig. 3A). As STAT3 has been previously implicated in astrocytic inflammatory responses, we performed western blotting experiments to confirm phosphorylation (activation) of STAT3. We found that phosphorylated STAT3 levels were significantly increased within 30 minutes of AA/OA addition to astrocytes (Extended Data Fig. 3B-D), a timescale consistent with direct activation of this pathway^41,42^. STAT3-mediated transcription is associated with reactive astrogliosis and pro-inflammatory secretion^43–45^, and acetylation of STAT3 has previously been implicated as a driver of astrocytic neurotoxicity upon chronic OXPHOS depletion in a mouse model^8^. Consistent with STAT3 phosphorylation, GSEA results showed significant enrichment of genes involved in other inflammatory pathways including IL6/JAK/STAT3 signaling and the interferon alpha (IFN-α) and gamma (IFN-γ) responses (Fig. 1C).

We also noted that two of the transcription factors whose activity was enriched following AA/OA treatment, AP1 and CREB, can be activated by the Mitogen-Activated Protein Kinase (MAPK). This is a class of kinases activated in response to cellular stress to induce an adaptive state. We performed a GSEA and found that transcripts affiliated with the MAPK signaling pathway are enriched in WT astrocytes treated with AA/OA compared to those treated with vehicle control (Fig. 4B). Upon examination of specific transcripts most upregulated following AA/OA treatment in astrocytes, three of these MAPK-associated transcripts encode proteins associated with negative regulation of inflammatory pathways. Growth arrest and DNA damage-inducible 45 proteins (Gadd45a/Gadd45b) are negative regulators of c-Jun Terminal Kinase (JNK), which is one of the MAPKs, and Zc3h12a limits NF-κB activation, again implicating a prominent NF-κB response to oxidative damage (Fig. 4C). Additionally, three more of these transcripts encode Vascular endothelial growth factor A (Vegfa), HGF receptor c-Met (Met), and Sphingosine kinase 1 (Sphk1), proteins that canonically promote cell survival (Fig. 4C). This suggested activation of kinases downstream of MAPK that canonically promote a cell survival response^46,47^.

To establish activation of p38/MAPK signaling as a component of the astrocytic response to mitochondrial damage, we performed western blots for the phosphorylated (active) form of p38. We found that p38 phosphorylation is significantly increased following 30 minutes of AA/OA treatment (Fig. 4D, E; Extended Data Fig. 3E-G). At that same timepoint, we also observed phosphorylation of both isoforms of JNK, p46 and p54, indicating the simultaneous activation of several MAPKs (Fig. 4D, F, G). Another of the five transcription factors whose activity is enriched in WT astrocytes post-damage, AP1 (Fig. 4A), is classically downstream of JNK signaling^48^. Although both p38/MAPK and JNK/SAPK are potentially damaging when active for an extended period, concurrent activation of NF-κB signaling is established to result in increased transcription of proteins that promote cell survival, as well as proteins that negatively regulate MAPK signaling to prevent prolonged activation and eventual apoptosis^49,50^.

Activation of p38/MAPK can be associated with pro-inflammatory or pro-survival transcriptional responses depending on the downstream signaling that ensues. We therefore sought to determine which kinases downstream of p38 are activated by astrocytic mitochondrial damage. Mitogen Activated Protein Kinase-Activated Protein Kinase 2 (MK2, also known as MAPKAPK2) and Mitogen Stress Kinases 1 and 2 (MSK1/2) are established downstream targets of p38. Cytoplasmic translocation of MK2 is facilitated by phosphorylation at T334 and is sometimes associated with stabilization of pro-inflammatory transcripts^51,52^. We detected expression of MK2 in astrocytes (Fig. 4D, Extended Data Fig. 3H), but we were unable to detect T334 phosphorylation following 30, 60, or 90 minutes of OXPHOS inhibitor treatment (Fig. 4H). Alternatively, p38/MAPK phosphorylation can activate Mitogen- and Stress-activated Kinases 1 and 2 (MSK1/2), kinases that reside upstream of CREB in the canonical p38/MAPK pro-survival signaling pathway^46,47^. We detected a ∼12-fold increase in MSK1 phosphorylation at T581 following 30 minutes of mitochondrial damage compared to baseline (Fig. 4D, I; Extended Data Fig. 3I, J).

CREB is among the transcription factors identified via GSEA as being active in astrocytes following OXPHOS inhibition (Fig. 4A) and is also downstream of MSK1 activation. We thus tested whether CREB is phosphorylated following mitochondrial damage in astrocytes. We observed significantly increased levels of phosphorylated CREB 30, 60, and 90 minutes following AA/OA treatment (Fig. 4J). These results indicate that astrocytes mount a canonically pro-survival p38/MAPK-facilitated response to mitochondrial damage.

### Loss of PINK1 or Parkin differentially alters inflammatory responses to mitochondrial damage in astrocytes

PINK1 and Parkin are key to mitochondrial damage response pathways in neurons, and loss-of-function mutations in either gene are causative for PD^53^. Thus, we, we performed bulk RNA-seq on astrocytes isolated from PINK1^-/-^ and Parkin^-/-^ mice treated with OXPHOS inhibitors, in parallel with our transcriptomic analysis of the mitochondrial damage response in WT astrocytes (Fig. 5A). PCA analysis of the resulting transcriptomic data from either PINK1^-/-^ or Parkin^-/-^ astrocytes post-mitochondrial damage showed separation from their respective controls along the first principal component, much like we observed for WT astrocytes (Fig. 5B). Further, the stressed astrocytes grouped together along the first principal component regardless of genotype, suggesting that the mitochondrial stress response in astrocytes is robust enough to account for the majority of the variability across the three genotypes tested (Extended Data Fig. 4A, B). Although AA/OA stress accounted for the majority of changes observed, we also observed differences in gene regulation at baseline and in response to mitochondrial damage across these different genotypes (Fig. 5C, D; Extended Data Fig. 4C-E).

**Figure 5.**
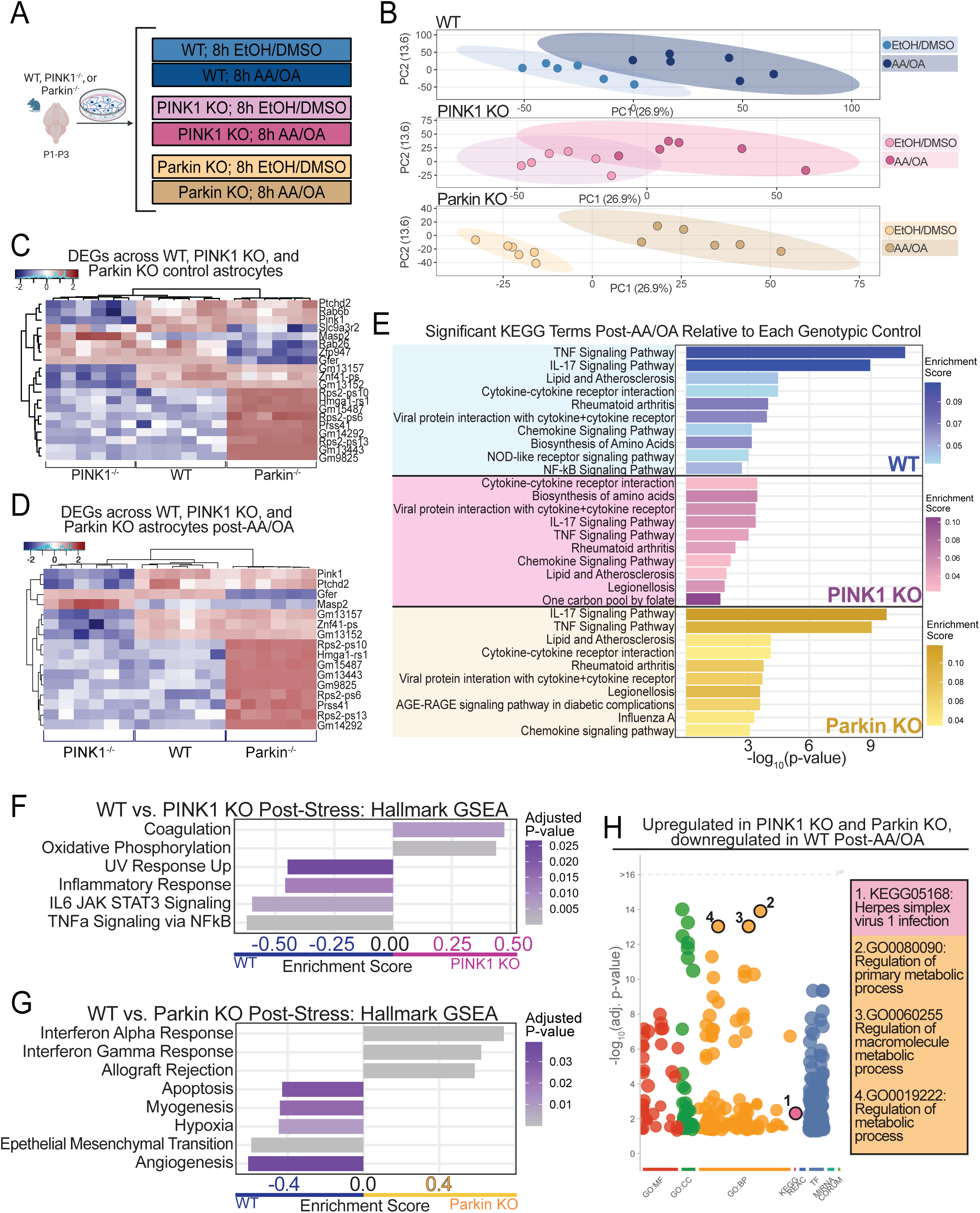
Loss of PINK1 or Parkin differentially alters inflammatory responses to mitochondrial damage in astrocytes. A) Schematic of bulk RNA sequencing experimental design. B) Principal component analysis depicting each of the biological replicates for each genotype, treated with either 10 μM AA/ 5 μM OA or vehicle control for 8h. Ellipses indicate 95% confidence regions based on a multivariate t-distribution (n=6 for each of the six conditions). C, D) Heat map of top differentially expressed gene modules between genotypes for vehicle control-treated astrocytes (top) and 8h 10μM AA/ 5μM OA treated-astrocytes (bottom). Adjacent phylograms illustrate hierarchal clustering results. Expression values are row-scaled to display relative expression across different samples (n=6 for each of the six conditions). E) Top 10 most significantly enriched KEGG terms as determined by GO analysis for each genotype post-AA/OA relative to the respective genotypic vehicle control (n=6 for each of the six conditions). F) GSEA results for WT astrocytes following 8h treatment with AA/OA compared to PINK1^-/-^ astrocytes following 8h treatment with 10 μM AA/ 5 μM OA, highlighting the differentially enriched Hallmark gene set terms (n=6 for both WT post-AA/OA and PINK1^-/-^ post-AA/OA). G) GSEA results for WT astrocytes following 8h treatment with AA/OA compared to Parkin^-/-^ astrocytes following 8h treatment with 10 μM AA/ 5 μM OA, highlighting the differentially enriched Hallmark gene set terms (n=6 for both WT post-AA/OA and Parkin^-/-^ post-AA/OA). H) GO analysis of genes differentially regulated across genotypes following 8h treatment with 10 μM AA/ 5 μM OA. Genes were included in the analysis if they were upregulated in both PINK1^-/-^ and Parkin^-/-^astrocytes (log_2_FC > 0), but downregulated or unchanged in WT astrocytes (log_2_FC ≤ 0). FDR correction was applied. Biological replicates are defined as independent experiments performed on astrocytes or neurons from different dissections.

Upon comparing AA/OA-treated astrocytes to each of their genotypic controls, we found that TNF-α signaling was not the most significantly enriched KEGG term following AA/OA treatment in PINK1^-/-^ or Parkin^-/-^ astrocytes as compared to their respective baselines (Fig. 5E). Further, we observed the appearance of several terms not present in the 10 highest enrichment scores for the KEGG analysis of WT astrocytes. This includes “legionellosis” in PINK1^-/-^ astrocytes following damage, indicative of a cellular response similar to that which is classically mounted following bacterial infection, and “Influenza A” in the Parkin^-/-^ astrocytes, indicative of a viral immune response. To determine how loss of either PINK1 or Parkin influences astrocytic responses to mitochondrial damage, we next directly compared the WT astrocyte transcriptome following AA/OA treatment to either PINK1^-/-^ or Parkin^-/-^ astrocyte transcriptomes following AA/OA treatment. We found that both TNF-α signaling via NF-κB and IL6/JAK/STAT3 signaling are significantly enriched in WT astrocytes following AA/OA treatment compared to PINK1^-/-^ astrocytes following AA/OA treatment (Fig. 5F). Moreover, AA/OA treatment in PINK1^-/-^ astrocytes failed to elicit a significant increase in TNF-α transcription as detectable by qPCR (Extended Data Fig. 4F). This demonstrates that PINK1 contributes substantially to the astrocytic immune response to mitochondrial damage in addition to initiating mitophagy. This could be explained via a mechanism previously identified by our group in which PINK1 and Parkin mediate NF-κB activation via recruitment of the NF-κB effector molecule NEMO to the ubiquitin chains formed on damaged mitochondria during mitophagy initiation^38^. One of the proteins most upregulated in PINK1^-/-^ astrocytes relative to both WT and Parkin^-/-^ astrocytes post-AA/OA treatment is Mannan-binding lectin-associated serine protease 2 (MASP2), which promotes activation of the complement immunity pathway^54,55^ (Fig. 5C, D, Extended Data Fig. 4G). This further suggests alternative pro-inflammatory signaling in PINK1^-/-^ astrocytes post-damage. In Parkin^-/-^ astrocytes, we observed an enrichment in transcriptomic signatures indicative of interferon responses following mitochondrial damage when compared to the WT response to the same damage stimulus (Fig. 5G). Specifically, both interferon alpha (IFN-α) and interferon gamma (IFN-γ) responses were enriched in Parkin^-/-^ astrocytes. Given that IFN-α-mediated signaling is predominantly associated with the leukocyte viral response and IFN-γ-mediated signaling is involved in macrophage activation in response to viral or bacterial invasion, this indicates that Parkin^-/-^ astrocytes also have a mitochondrial damage response that involves enhanced pro-inflammatory signaling distinct from that of the NF-κB-mediated response in WT astrocytes.

We also observed an enriched hypoxia response in WT astrocytes post-AA/OA compared to Parkin^-/-^astrocytes (Fig. 5G). Of several previously described astrocytic reactive states, the “A2, neuroprotective” state is induced by hypoxic insult. We thus measured expression of these “A2” astrocyte markers in our dataset and observed enrichment of these genes in astrocytes post-AA/OA treatment (Extended Data Fig. 4H), suggesting that an A2 state transition could be driven by OXPHOS failure. This enrichment post-AA/OA was not observed for “A1, pro-inflammatory” astrocyte markers (Extended Data Fig. 4I) or pan-reactive astrocyte markers (Extended Data Fig. 4J). Moreover, the most substantial enrichment of A2 markers was in WT astrocytes (Extended Data Fig. 4H). While the A2 reactive state is described as anti-inflammatory, recent work suggests that persistent A2 astrocytes can evolve to take on an A1-like, pro-inflammatory response^56^, emphasizing the necessity of negative feedback mechanisms such as accelerated clearance of damaged mitochondria that could resolve the signaling driving this physiological change.

To interrogate how loss of mitophagy might impact the astrocytic response to mitochondrial damage, we extracted the differentially expressed genes that were significantly upregulated in both PINK1^-/-^ and Parkin^-/-^ astrocytes post AA-OA treatment and downregulated in WT astrocytes post-AA/OA treatment and performed a GO analysis on these genes (Fig. 5H, Extended Data Fig. 4K). We observed upregulation of various metabolic processes, which could potentially represent activation of compensatory networks triggered in the face of irresolvable mitochondrial dysfunction. We found that the only KEGG term significantly enriched in this set of genes was Herpes Simplex Virus 1, again indicating an exacerbated viral-type inflammatory response to mitochondrial damage when mitophagy is lost. Altogether, these results indicate that the loss-of-function mutations in PINK1 and Parkin identified in some familial Parkinson’s disease patients may manifest in altered astrocytic immune signaling following accumulation of damaged mitochondria.

### Human astrocytes from the substantia nigra of PD-affected tissue show enrichment of the same inflammatory pathways as AA/OA-treated astrocytes *in vitro*

Previous work has demonstrated that mitochondrial damage occurs in astrocytes during normal aging and in astrocytes from patients with Parkinson’s disease (PD)^28–30,57^. To determine whether the inflammatory signatures we identified in WT murine astrocytes could also be seen in human disease, we accessed single cell RNA-sequencing data for human brain tissue from either Parkinson’s-positive samples or age-matched controls (courtesy of the Michael J. Fox Foundation’s Aligning Science Across Parkinson’s initiative)^58^ (Extended Data Fig. 5A, B). We isolated data from cells annotated as astrocytes harvested from the substantia nigra and performed a clustering analysis using Seurat. This analysis revealed distinct transcriptomic clusters in a UMAP-reduced space (Fig. 6A). To validate that the cells analyzed were astrocytes, we looked for enrichment of protoplasmic and fibrous astrocyte markers (Extended Data Fig. 6A, B). We found only one transcriptomic cluster that was not enriched for astrocyte markers, and instead showed enrichment for markers of oligodendrocyte lineage cells (Extended Data Fig. 6A, B); we excluded this cluster when performing further analyses. Interestingly, we noted that some transcriptomic clusters consisted almost exclusively of astrocytes from PD-positive samples (cluster 10), and others were comprised almost exclusively of astrocytes derived from age-matched control samples (cluster 6; Fig. 6B, Extended Data Fig. 5C, D). Specifically, the PD-enriched cluster was comprised of >99% cells from PD-positive samples, and the control-enriched cluster was comprised of 96% cells derived from control samples (Fig. 6B). This suggests that distinct populations of astrocytes emerge through normal aging as compared to aging with PD.

**Figure 6.**
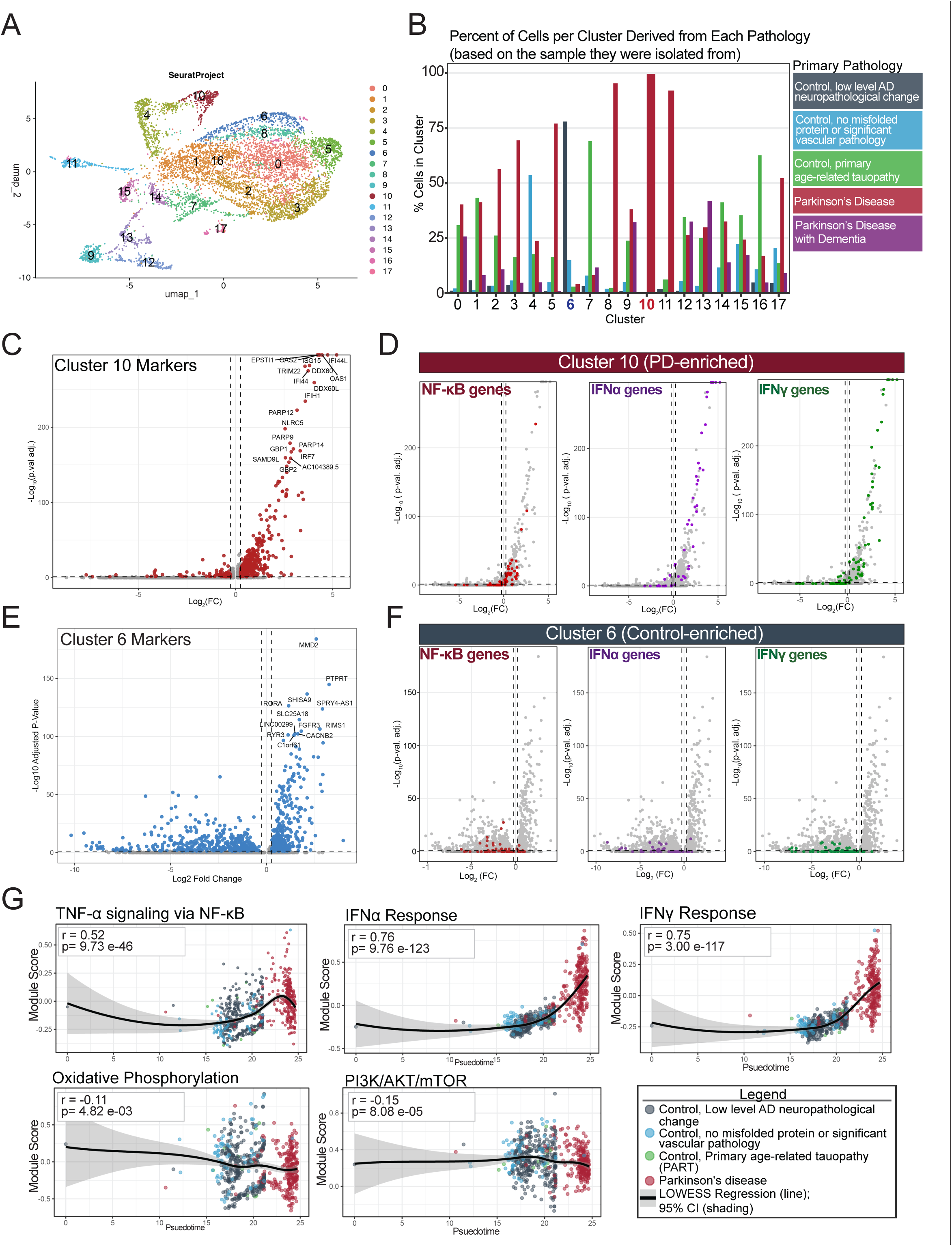
Human astrocytes from the substantia nigra of PD-affected tissue show enrichment of the same inflammatory pathways as AA/OA-treated astrocytes in vitro. A) Seurat-generated transcriptomic clusters of astrocytes isolated from the substantia nigra of control or PD+ human brain tissue. B) Cluster composition colorized by the percentage of cells in each cluster that are derived from samples designated as being each of the five possible pathologies. The vast majority of cells in cluster 6 are derived from control samples, whereas the vast majority of cells in cluster 10 are derived from PD+ samples. C) Gene markers for cluster 10 (PD-enriched). Markers with a | Log_2_(FC) | > 0.25 and an adj. p-value <0.05 are highlighted (red). D) Markers for cluster 10 with genes belonging to the TNF-α signaling via NF-κB (red), IFN-α response (purple), and IFN-γ response (green) Hallmark pathways highlighted. Lines at | Log_2_(FC) | > 0.25 and adj. p-value <0.05. E) Gene markers for cluster 6 (Control-enriched). Markers with a | Log_2_(FC) | > 0.25 and an adj. p-value <0.05 are highlighted (blue). F) Markers for cluster 6 with genes belonging to the TNF-α signaling via NF-κB (red), IFN-α response (purple), and IFN-γ response (green) Hallmark pathways highlighted. Lines at | Log_2_(FC) | > 0.25 and adj. p-value <0.05. G) Module scores across a psuedotime trajectory from the control to the PD-enriched cluster for five Hallmark pathways of interest. Dots represent individual cells colorized by the pathology classification of the sample they are derived from. Biological replicates are defined as independent experiments performed on astrocytes or neurons from different dissections. For all graphs, p < 0.05 = *; p < 0.01 = **.

To determine whether signatures of mitochondrial damage observed in our analysis of RNA-seq data from astrocytes *in vitro* treated with OXPHOS inhibitors are present in astrocytes isolated from PD-positive samples, we further characterized the control- and PD-enriched clusters. In our *in vitro* astrocyte model, we observed significant enrichment of transcripts associated with TNF-α signaling via NF-κB in WT astrocytes following AA/OA treatment. Further, transcripts affiliated with the IFN-γ and IFN-α responses were significantly enriched in Parkin^-/-^ astrocytes when compared to WT astrocytes post-damage. To determine how this compares to the human astrocyte data, we quantified the number of transcriptomic markers for our control- and PD-enriched clusters that are classified as being a part of the NF-κB, IFN-α, or IFN-γ pathways (Fig. 6C-F). Of the transcripts significantly upregulated in the PD-enriched cluster, 27 are implicated in NF-κB signaling, 30 in IFN-α signaling, and 43 in IFN-γ signaling. Of the transcripts significantly downregulated in the PD-enriched cluster, 3 are implicated in NF-κB signaling, 2 in IFN-α signaling, and 5 in IFN-γ signaling. Conversely, of the transcripts significantly upregulated in the control-enriched cluster, only 2 are implicated in NF-κB signaling, 1 in IFN-α signaling, and 0 in IFN-γ signaling. Of the transcripts downregulated in this cluster, 16 are implicated in NF-κB signaling, 13 in IFN-α signaling, and 18 in IFN-γ signaling. These data illustrate that transcripts associated with NF-κB activation, IFN-α signaling, and IFN-γ signaling were generally upregulated the PD-enriched cluster and downregulated the control cluster, emphasizing the relevance of these pathways in PD.

We next calculated a pseudotime trajectory from the control-enriched cluster to the PD-enriched cluster to further investigate transcriptomic differences between these groups (Extended Data Fig. 5E). We iteratively calculated the correlation of each of the 50 hallmark pathway module scores with this pseudotime trajectory (Extended Data Fig. 5F). We found that the hallmark pathways associated with IFN-γ and IFN-α bore the strongest correlation with graded transcriptomic changes across pseudotime (Fig. 6G). This is corroborated by GSEA analysis conducted on transcripts that are significantly upregulated in the PD-enriched cluster compared to the control-enriched cluster (Extended Data Fig. 5G), and is congruent with our data indicating that Parkin^-/-^ astrocytes following AA/OA treatment are significantly enriched for IFN-α and IFN-γ response-associated transcripts compared to WT astrocytes following AA/OA treatment (Fig. 5G). Among the pathways negatively correlated with the transition from the control-enriched to the PD-enriched cluster were oxidative phosphorylation, corroborating prior studies using PD tissue that suggest astrocytes undergo OXPHOS impairments in PD, and PI3K/Akt/mTOR signaling, which relates to autophagic maintenance of cellular contents such as organelles and proteins and is also thought to decline in efficiency in diseases of aging like PD. These data provide evidence for an astrocytic pro-inflammatory response present in PD that does not occur in healthy aging, and which resembles the transcriptomic changes induced by mitochondrial damage that were observed in our *in vitro* astrocyte model.

## DISCUSSION

Here we find that OXPHOS failure in astrocytes is sufficient to initiate inflammatory signaling modulated by PINK1 and Parkin, and that the resulting inflammatory signature is also evident in a transcriptomic analysis of astrocytes derived from postmortem tissue of patients with PD (Fig. 7). It is well-established that astrocytic mitochondrial health declines with age^5,28,29^; this decline is exacerbated in PD^30^, emphasizing the importance of understanding how mitochondrial dysfunction influences astrocyte physiology. We now identify a self-limiting pathway in which OXPHOS failure induces PINK1/Parkin-dependent mitophagy, while simultaneously leading to induction of NF-κB signaling in primary astrocytes. NF-κB activation in turn induces the upregulation of the autophagy receptor p62 and enhanced mitochondrial turnover. This negative feedback loop provides a mechanism by which transient NF-κB activation can protect against chronic inflammation in the nervous system in response to mitochondrial injury. Mechanisms limiting inflammatory signaling in astrocytes are especially important, as they directly influence neuronal health and function^59,60^.

**Fig. 7.**
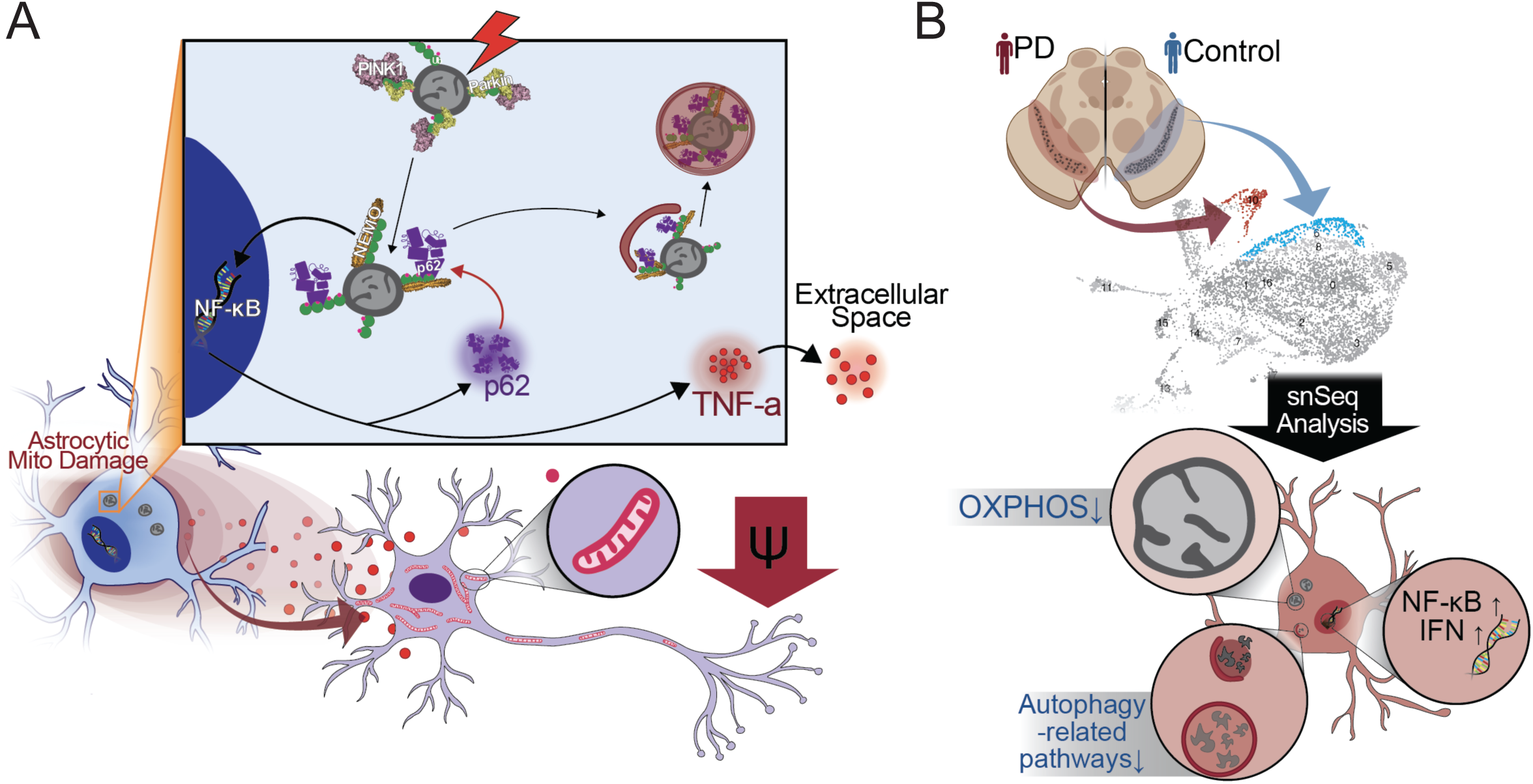
Summary. A) Mitochondrial damage in astrocytes activates NF-κB signaling, which leads to secretion of TNF-α and heightened efficiency of mitochondrial turnover. The same insult to PINK1^-/-^ or Parkin^-/-^ astrocyte models of PD causes alternative inflammatory signaling that parallels what we observe in transcriptomic analysis of human postmortem tissue from PD patients (B).

Consistent with our model, temporally limiting astrocyte-specific NF-κB activation in an Alzheimer’s disease mouse model has been found to be pivotal for neuronal survival; induction of transient and astrocyte-specific NF-κB activation facilitates a protective response to protein aggregation, whereas chronic NF-κB activation in astrocytes is sufficient to drive neuronal senescence and cause inflammation in the hippocampus^61^. NF-κB activation is a recognized aspect of PD pathology as well, though its pathological role has not been clear^62,63^. In dopaminergic neurons in postmortem PD tissue, the major transcription factor mediating NF-κB signaling is more than 70-fold enriched in nuclei compared to control tissue^63^, suggesting chronic activation of NF-κB signaling in late-stage disease. Thus, we postulate that the transient NF-κB activation induced by acute OXPHOS inhibition encourages clearance of damaged mitochondria, preventing activation of multiple pro-inflammatory pathways. However, a failure to remove damaged mitochondria and inactivate NF-κB signaling may cause astrocytes to drive neurotoxicity.

Our data suggest that the the astrocytic response to mitochondrial damage is both pro-inflammatory and pro-survival, our work shows that NF-κB activation following OXPHOS inhibition in astrocytes induces the transcriptional changes necessary to mitigate mitochondrial damage and facilitate recruitment of microglia to limit the spread of acute injury. Our data also demonstrate that these responses are altered in astrocytes lacking PINK1 or Parkin. In contrast with WT astrocytes, the inflammatory response mounted by PINK1^-/-^astrocytes following mitochondrial damage was significantly less enriched for transcriptional signatures of NF-κB activation. Previous work from our lab has shown that astrocytes recruit NEMO to ubiquitin chains constructed on OMM proteins in a PINK1/Parkin-dependent manner^38^. Eliminating this pathway for NF-κB activation is sufficient to alter the inflammatory landscape of these cells, preventing transient upregulation of genes that promote survival and instead promoting upregulation of alternative inflammatory proteins such as Masp2 that may not be intrinsically self-limiting and could instead lead to neurotoxicity^54,55^. Parkin-null cells exhibit a distinct proinflammatory response. Our transcriptomic analysis shows that the response to OXPHOS failure in Parkin^-/-^ astrocytes is enriched for IFN-α and IFN-γ response signatures compared to WT astrocytes, indicating activation of alternative inflammatory signaling pathways also implicated in PD pathology^62^. Active IFN-γ signaling can be amplified by coincident TNF-α signaling, as shown in peritoneal macrophages^64,65^, suggesting that co-activation of NF-κB and IFN-γ could promote chronic inflammatory signaling. These *in vitro* data, and the observed changes in astrocytic cytokine and chemokine secretion following OXPHOS inhibition, identify persistent mitochondrial damage in astrocytes as a potential source of NF-κB and interferon signaling in the brain.

In line with our in vitro findings, our analysis of human postmortem single cell sequencing data identified activation of NF-κB, IFN-γ, and IFN-α in a transcriptomic cluster of astrocytes from the substantia nigra that was enriched in PD patients, suggesting that all three pathways are activated during disease. Pseudotime analysis indicates that IFN-γ and IFN-α were most closely correlated with graded transcriptomic changes between control and patient datasets, further implicating these inflammatory pathways in disease progression.

To date, most research on transcriptomic changes in astrocytes has focused on how these cells respond to extracellular cues^66–69^. There has been particular interest in astrocytic responses to compounds (C1q, TNF-α, and IL-1α) secreted by microglia that prompt a transition to a well-defined, pro-inflammatory astrocytic state^59,66^. These foundational studies helped shape our understanding of astrocytes as environmentally responsive and transcriptionally adaptive cells that directly influence neuronal health. Here, we build on this body of work by examining the consequences of direct damage to astrocytes via OXPHOS inhibition. Our work defines downstream signaling responses such as TNF-α release that directly impact neuronal health. We observed pro-inflammatory shifts, mediated in part by activation of NF-κB signaling in response to mitochondrial damage. However, we also see enrichment of a canonically pro-survival arm of the MAPK pathway^70–74^ and “A2, anti-inflammatory” reactive astrocyte markers that were originally described in response to middle cerebral artery occlusion in mice^59^. This supports the model that there is a spectrum of profiles that astrocytes adopt in a stressor- and timing-dependent manner^75^, and reinforces our hypothesis that there are both pro-inflammatory and pro-survival facets of the astrocyte mitochondrial damage response. While the pro-inflammatory astrocyte transition that occurs in response to environmental cues is understood to be facilitated by synthesis of long-chain fatty acids^76^, the A2 astrocyte transition characterized in that same initial work demonstrating reactive astrogliosis has not been mechanistically defined. Our results demonstrate that loss of OXPHOS, a plausible consequence of the hypoxic injury that elicits this A2 astrocyte transition in mice, is sufficient to induce upregulation of A2 genes *in vitro*. Although the A2 state was originally defined as anti-inflammatory due to its initially beneficial impact on neuronal health^59^, more recent work has shown that A2-like astrocytes can eventually transition to a more pro-inflammatory, neurotoxic state^56^. This highlights the need for built-in negative feedback mechanisms such as the one we identify to prevent prolonged and potentially neurotoxic signaling^77^. We thus propose that astrocyte-autonomous responses to internal stressors, much like their responses to factors secreted by pro-inflammatory microglia, influence astrocytic signaling in a manner that has direct consequences for neuronal health.

There are several important questions that remain unanswered. Regional differences in astrocyte transcriptomes *in vivo*^78,79^ might predispose certain subgroups of astrocytes to a heightened or altered response to OXPHOS failure. Further, cytokine profiling detected secretion of compounds in response to OXPHOS inhibition in astrocytes that microglia are well-established to detect, suggesting that NF-κB-dependent secretion following mitochondrial damage in astrocytes may be sufficient to elicit an inflammatory response in microglia^80–82^. Further work will be required to address these complex intercellular relationships.

In summary, our data demonstrate that astrocyte-autonomous OXPHOS failure initiates PINK1/Parkin-dependent mitophagy, and in parallel induces a transcriptional program in which NF-κB activation is central. This transcriptional response promotes accelerated mitochondrial turnover, serving as a negative feedback loop to limit pro-inflammatory signaling. These transcriptomic changes also lead to NF-κB-dependent changes in cytokine secretion that likely impact both neurons and microglia. Loss of either PINK1 or Parkin inhibits mitophagy and leads to differential activation of inflammatory cascades that may not be self-limiting. Grounding the physiological relevance of our findings, we identified activation of NF-κB and interferon signaling pathways in our analysis of PD-derived human astrocyte transcriptomes. Together, these data establish the impact of acute mitochondrial damage in astrocytes. Astrocytes have long been viewed as a conduit for microglial-instigated neuroinflammation rather than an autonomous source of pro-inflammatory stimuli in the context of neurodegenerative disease; our findings extend the current paradigm to include the contributions of astrocyte-autonomous sources of inflammatory signaling, which may prove especially important for treating diseases like PD where neuroinflammation is a key pathological hallmark.

## Acknowledgements

We thank Kaya J.E. Matson for her input regarding RNA-sequencing and analysis, Karen Jahn for breeding and maintaining the mice used in this study, Mariko Tokito for her technical assistance. We are also grateful to the Korb laboratory at the University of Pennsylvania for sharing their library generation protocol, and the Penn Next Generation Sequencing Core for sequencing services.

We are grateful to the Aligning Science Across Parkinson’s (ASAP) team Hurley, especially coordinator Dorotea Fracchiolla, for providing insights throughout the course of this work. Single cell sequencing data used in the preparation of this article was obtained from the following collections from the Aligning Science Across Parkinson’s Collaborative Research Network Cloud (ASAP CRN Cloud) (RRID:SCR_023923): Postmortem-derived Brain Sequencing Single Cell (sc)RNAseq Collection (cohort-pmdbs-sc-rnaseq, v2.0.0; DOI: https://doi.org/10.5281/zenodo.14270014. A full list of contributors to the ASAP CRN Cloud is available at https://bit.ly/CRN_Cloud_Authorship_List. We would like to specifically acknowledge the Jakobssen, Lee, Hafler, Hardey, and Scherzer ASAP teams, who generated the harmonized single cell sequencing dataset repurposed for analysis in this study.

This work was funded by an Aligning Science Across Parkinson’s grant ASAP-000350 through the Michael J. Fox Foundation for Parkinson’s Research (E.L.F.H), an NSF Graduate Research Fellowship DGE-2236662 (J.F.R.), an NIH National Research Service Award 1F31NS143320-01 through the National Institute of Neurological Disorders and Stroke (J.F.R.), and a Penn First Exposure to Biomedical Research Fellowship (C.V.R.).

## Author information

### Contributions

J.F.R and E.L.F.H conceptualized the study, with C.V.R providing input. J.F.R and C.V.R performed experiments and data analysis. J.F.R wrote the original draft of the paper, and J.F.R and E.L.F.H reviewed and edited the paper.

## METHODS

### Ethics statement

The work detailed in this manuscript was performed in accordance with the Guide for the Care and Use of Laboratory Animals of the National Institutes of Health (NIH). All experiments using animals followed protocol approved by the Institutional Animal Care and Use Committee (IACUC) at the University of Pennsylvania.

### Cell culture

#### Astrocyte monoculture and transfection

Astrocytes were derived from murine models, and the WT model used is the recommended genetic control for both the PINK1 and Parkin knockout mice (WT, C57BL/6J; PINK1^-/-^, B6.129S4-Pink1^tm1Shn^/J; Parkin^-/-^, B6.129S4-Prkn^tm1Shn^/J). Astrocyte-enriched cultures (90-95%) were generated as previously described^83^. Briefly, murine cortices were isolated from mice on postnatal days 1-3, digested in trypsin (0.25%), and plated on tissue culture treated flasks coated with poly-d-lysine (ThermoFisher). Cells were grown for two weeks in maintenance media (DMEM with high glucose, L-glutamine, pyruvate, and without HEPES [Gibco 11995065] with 10% FBS [Cytiva] and 50 U/ml penicillin and 50 mg/ml streptomycin) and placed on a shaker at 37C, 250 rpm for ∼6 hours. Cells were then washed twice with PBS (pH 7.4, no calcium, no magnesium [Gibco]) and dissociated from the plate using trypsin with EDTA. Trypsin was neutralized with maintenance media and cells were pelleted. Pelleted cells were washed thrice with maintenance media and resuspended via trituration using a transfer pipette followed by a p1000. Prior to plating, cells were passed through a sterile, 70 μm-pore cell culture filter to remove any debris. Astrocytes were then plated on PDL-coated plates and treated with the mitotic inhibitor cytosine arabinoside (AraC; final concentration 1 μM) once adherent to prevent overgrowth. Astrocytes were used for experiments between 10-14 days post-passage. To visualize YFP-Parkin or a YFP control, astrocytes were transfected for 45 minutes with 0.75 μg of plasmid using lipofectamine 2000 per the manufacturer’s protocol 24-48 hours prior to treatment and fixation.

#### Rat hippocampal neuron culture and transfection

Sprague–Dawley rat hippocampal neurons (harvested at embryonic day 18) were obtained from the Neurons R Us Culture Service Center at the University of Pennsylvania (RRID: SCR_022421). Cells (200k/dish) were plated in glass-bottom 35-mm dishes (20mm glass; MatTek) precoated with 0.5mg/ml poly-L-lysine (Sigma-Aldrich). Neurons were plated in attachment media (MEM [Gibco], 10% horse serum, 33 mM D-glucose, 1 mM sodium pyruvate). Once neurons were adherent, attachment media was replaced with maintenance media (neurobasal [Gibco] with 33 mM D-glucose [Sigma-Aldrich], 2 mM GlutaMAX [Invitrogen], 100 U/ml penicillin and 100 mg/ml streptomycin [Gibco], and 2% B-27 5x supplement [Thermo Fisher Scientific]). Neurons were maintained at 37°C in 5% CO_2_; AraC (final concentration 1 µM) was added 24 hours after plating. Half the media was replaced with fresh maintenance media on day *in vitro* (DIV) 7. Neurons were transfected with mito-mEmerald 24-48 hours before use with lipofectamine 2000 as per the manufacturer’s protocol and live imaged on DIV 14.

#### iPSC-derived neuron (iNeuron) generation and iNeuron/astrocyte co-culture

KOLF2.1J-background WT iPSCs^84^ were gifted to our lab by B. Skarnes (Jackson Laboratory for Genomic Medicine, Farmington, CT, USA) as part of the iPSC Neurodegenerative Disease Initiative (iNDI) and were validated by karyotyping and mycoplasma testing as described^85^. Doxycycline-inducible hNGN2 was delivered using PiggyBac. Thawed cells were cultured on plates coated with Growth Factor Reduced Matrigel (Corning) and fed daily with Essential 8 media (Thermo Fisher Scientific). After 72h, transfected iPSCs were selected for 48h with 0.5 μg/ml puromycin (Takara). Differentiation into iNeurons was performed using an established protocol^84,86^ in which iPSCs were passaged using Accutase (Sigma-Aldrich) and plated on Matrigel-coated dishes in Induction Media (DMEM/F12 supplemented with 1% N2-supplement [Gibco], 1% NEAA [Gibco], 1% GlutaMAX [Gibco], and 2 μg/ml doxycycline). After 72 h of doxycycline-induced differentiation, iNeurons were dissociated with Accutase and cryo-preserved in liquid nitrogen. For use in co-culture, pre-differentiated iNeurons were thawed, plated on glass-bottomed dishes coated with poly-L-ornithine (175k iNeurons/dish) and cultured for seven days in iNeuron media (BrainPhys Neuronal Media [StemCell] supplemented with 2% B-27 [Gibco], 10 ng/ml BDNF [PeproTech], 10 ng/ml NT-3 [PeproTech], and 1 μg/ml laminin [Corning]). On day seven, iNeuron media was replaced with the same media containing an additional supplement, G-5 Supplement (1%; Gibco). Murine astrocytes (10-14 days post-passage) were dissociated using Trypsin/EDTA and resuspended in iNeuron media with 1% G-5 supplement. Astrocytes (65k/dish) were added to neurons.

Astrocytes (1 million) were transfected with 1 μg human YFP-Parkin using lipofectamine 2000 transfection reagent as per the manufacturer’s instruction. After 24 hours, transfected astrocytes were resuspended in iNeuron media with G-5 supplement (1%; Gibco) and added to iNeurons. Transfected astrocytes (65k/dish) and iNeurons (175k/dish) were grown in co-culture for six days until use to allow ample time for astrocyte branching.

#### Small molecule inhibitor treatments

To induce mitochondrial damage via OXPHOS failure, cells were co-treated with Antimycin A (10 μM unless otherwise noted; Sigma Aldrich A8674) and oligomycin A (5 μM unless otherwise noted; Sigma Aldrich 75351). Imaging experiments were performed 2h post-OXPHOS inhibitor treatment unless otherwise noted. To inhibit NF-κB-mediated signaling, astrocytes were pre-treated with 5 μM BMS-345541 (Abcam) for the 48 hours leading up to the experimental treatment, with fresh media and inhibitor being added every 24h. The same concentration of inhibitor was then used for the duration of the experimental treatment. To block lysosome acidification, cells were treated with 10nM bafilomycin A1 (Sigma) concurrently with the indicated experimental treatment.

### RNA-sequencing and analysis

#### cDNA library preparation

To generate cDNA libraries, 2.75 million astrocytes were harvested and RNA was isolated using the standard TRIzol protocol. RNA was further purified using an RNA clean and concentrator kit (Zymo Reseach R1017) in conjunction with the recommended DNase I digestion step (Zymo E1010). Libraries were generated using the TruSeq Stranded mRNA kit as directed. The generated cDNA libraries were cleaned using AMPure XP beads (Beckman, A63880) both prior to adenylation and after library generation. The batch in which libraries were prepared had no bearing on sample variability (Extended Data Fig. 4B).

Library quality control checks, pooling, and sequencing were performed by the Next Generation Sequencing Core at the University of Pennsylvania (RRID:SCR_024999). Library quality was confirmed using a bioanalyzer, and library concentrations were determined using the NEBNext Library Quant Kit for Illumina (New England Biolabs) to ensure equimolar pooling prior to loading onto two lanes of an S1 flow cell. As there were six biological replicates total for each condition, three biological replicates from each condition were loaded into each lane. The flow cell lane in which samples were run also did not influence sample variability based on principal component analysis. Paired-end, 100bp read length sequencing with a read depth of at least 30 million per sample was performed using a NovaSeq X Plus (Illumina).

#### Bulk RNA-sequencing data analysis

Bulk RNA-sequencing data was processed and analyzed using a previously established workflow as a framework^87^. Briefly, reads were reads were pseudoaligned to a reference transcriptome derived from the Ensembl genome build GRCm39 using Kallisto. Data were imported into RStudio and gene identifiers were annotated using the EnsDb.Mmusculus.v79 package. Raw count data were filtered to remove lowly expressed genes prior to differential expression analysis. Specifically, genes with counts per million (CPM) greater than 1 in at least six samples were retained to ensure sufficient expression across samples to perform further statistical testing. Filtered count data were normalized to account for library size and composition using the trimmed mean of M-values (TMM) method, which was implemented though the edgeR package.

To identify genes upregulated post-stress relative to the control for each genotype, differentially expressed genes between stressed and control conditions were identified using the limma package with empirical Bayes moderation. Genes with a Benjamini–Hochberg–adjusted p-value < 0.01 and an absolute log_2_(fold change) of greater than or equal to 2 were selected for visualization. Heatmaps were generated using normalized expression values, and expression was scaled by row. Unsupervised hierarchical clustering was performed on genes using Pearson correlation and on samples using Spearman correlation, with complete linkage. Gene modules to use for further analysis (GO analyses, GSEA analyses) were extracted by cutting the gene dendrogram for the two conditions being compared into two clusters (k=2).

Gene Ontology (GO) enrichment analysis was performed using the gprofiler2 R package. Differentially expressed genes were identified as described. The resulting gene list was used as input for GO enrichment analysis with the gost function. *Mus musculus* was specified as the reference organism and false discovery rate (FDR) correction was applied. Manhattan plots were generated using gostplot.

Gene set enrichment analysis (GSEA) was performed using the clusterProfiler R package. Hallmark gene sets for *Mus musculus* were obtained from the Molecular Signatures Database (MSigDB) using the msigdbr package. Genes were ranked by log_2_(fold change) for the indicated comparison between conditions. The ranked gene list was sorted in decreasing order of log_2_(fold change) and used as input for pre-ranked GSEA. Enrichment analysis was conducted using the GSEA function with default parameters. A fixed, random seed was set to ensure reproducibility. GSEA plots were visualized using the enrichplot package.

To determine transcription factors mediating astrocytic responses to mitochondrial damage, we ran a GSEA for terms in the TFT Legacy database of the MSigDB. We extracted the 50 most-enriched terms describing transcription factor activity and downloaded the lists of genes affiliated with each term. We then calculated the percent similarity between each list and every other list using the formula (number of genes in common/total number of genes) x 100), made a network depicting the similarities between these 50 terms using the qgraph package, and looked at the resulting network to determine which general classes of transcription factors were most represented in the 50 TFT-legacy terms with the highest positive enrichment scores. ATF and CREB were grouped together because they belong to the same family and had largely overlapping downstream targets as defined by the TFT Legacy database.

To identify pathways upregulated in PINK1^-/-^ astrocytes compared to WT astrocytes post-AA/OA, we iteratively performed GSEA analyses for each of the 50 hallmark pathways identified in the MSigDB comparing these two conditions and graphed pathways for which the adjusted p-value was <0.05. To identify DEGs upregulated in OXPHOS-deficient astrocytes without the ability to perform mitophagy, genes were selected that were upregulated in both PINK1^-/-^ and Parkin^-/-^ astrocytes (log2FC > 0) but downregulated or unchanged in WT astrocytes (log2FC ≤ 0). Enrichment analysis was performed using g:Profiler (mmusculus) with a false discovery rate correction. Manhattan plots were generated using the gostplot package.

MAPK-associated genes highlighted in the volcano plot in Fig. 3A are defined as genes in the GO BP term list (GO:1900745; GOBP_POSITIVE_REGULATION_OF_ P38MAPK_CASCADE). GSEA for p38/MAPK was run based on the positive regulation of p38MAPK cascade GO term (GO:1900745). TNF-α via NF-κB signaling-associated genes were highlighted based on the term list of the same name (GSEA: MM3860).

#### Single cell sequencing data analysis

Data used in the preparation of this article was obtained from the following collections from the Aligning Science Across Parkinson’s Collaborative Research Network Cloud (ASAP CRN Cloud) (RRID:SCR_023923). The dataset accessed was: Postmortem-derived Brain Sequencing Single Cell (sc)RNAseq Collection (cohort-pmdbs-sc-rnaseq, v2.0.0; DOI: https://doi.org/10.5281/zenodo.14270014). The data are controlled. Researchers can register for access to these data by submitting a Data Use Application through the ASAP CRN Cloud website (https://cloud.parkinsonsroadmap.org/collections). Data dictionaries, README files, protocols used to collect the data, and data processing pipelines are openly available at https://cloud.parkinsonsroadmap.org/collections. A full list of contributors to the ASAP CRN Cloud is available at https://bit.ly/CRN_Cloud_Authorship_List.^58^

For the purposes of this study, we accessed the integrated (harmonized) version of this data; as such, the data had already undergone mapping, processing, and batch correction and integration of all samples in the dataset as described on the platform where the data is accessible. As an overview for quality control, for which the full code is made available through the ASAP CRN platform, the authors of this dataset note in their description that the quality control filtering cutoffs were chosen to be as follows: mitochondrial gene percentage < 10%; doublet scores < 0.2; total counts between 500 and 100,000; and number features per cell between 300 and 10,000.

Of this dataset, we extracted the cells denoted as being astrocytes from the substantia nigra. This left us with cells from 41 different patient samples characterized as belonging to one of five classes depending upon the pathology the patient presented with: 1) Control, with a low level of Alzheimer’s disease neuropathological change; 2) Control, with no misfolded protein or significant vascular pathology; 3) Control, primary age-related tauopathy; 4) Parkinson’s disease; 5) Parkinson’s disease with dementia (Extended Data Fig. 5A). Of these 41 samples, only one sample derived from a patient with Parkinson’s was annotated as testing positive for a known familial mutation (GBA L444P). While astrocytes from this sample were included in the clustering analysis, none of these astrocytes were sorted into the clusters we defined as being control-or PD-enriched, so all analysis run on our two clusters of interest were samples from patients with sporadic PD. The ages at death for each sample are depicted in the panel regarding characterization of samples (Extended Data Fig. 5A).

Data for these cells annotated as being from the substantia nigra and identified as astrocytes was read into R as a Seurat object, and data were analyzed using the Seurat package following the general workflow recommended in the aforementioned DIY Transcriptomics course^87^. Briefly, highly variable genes were identified using a variance-stabilizing method, and the top 2,000 variable features were selected for downstream analysis. Principal component analysis (PCA) was performed on these variable genes. The first 15 principal components were used to construct a shared nearest neighbor (SNN) graph and to generate a Uniform Manifold Approximation and Projection (UMAP) embedding. The first 15 principal components were selected because that is where components began contributing diminishing variance based on visual inspection of an elbow plot. Cell clustering was performed using a graph-based Louvain algorithm with a resolution parameter of 0.5.

Pseudotime trajectory analysis was used to infer dynamic transitions between subpopulations of astrocytes. Cells sorted into clusters 6 (control-enriched) and 10 (PD-enriched) were extracted and converted to a SingleCellExperiment object, and a pseudotime trajectory was inferred beginning at the control-enriched cluster and terminating at the PD-enriched cluster using the Slingshot package. Pseudotime values were computed for each cell along this lineage and extracted using the slingPseudotime function to be mapped back onto the Seurat object for downstream analysis. To assess pathway-level changes along the inferred pseudotime trajectory, the expression of genes affiliated with each of the 50 Hallmark gene set pathways was quantified and correlated with pseudotime. Hallmark gene sets were obtained from MSigDB, and genes within each set were filtered to retain only those expressed in the dataset. For each pathway, a module score was calculated at the single-cell level using Seurat’s AddModuleScore function. Hallmark pathways with fewer than three genes present in the dataset were excluded from analysis. A Pearson’s correlation was calculated to assess relationships between module scores for individual pathways and the inferred pseudotime trajectory.

Differential gene expression analysis between clusters was performed using Seurat’s FindMarkers function. The control-enriched cluster and the PD-enriched cluster were compared bidirectionally to identify genes differentially expressed between the two populations. Genes were ranked by average log_2_(fold change) for downstream pathway analysis. Gene Set Enrichment Analysis (GSEA) was performed using the clusterProfiler package to identify biological pathways enriched between clusters. Ranked gene lists were generated based on average log_2_(fold change) values from the cluster comparison, with positive values indicating higher expression in cluster 10. Hallmark gene sets were obtained from the MSigDB database using the msigdbr package, and a GSEA for each hallmark pathway was conducted. Enrichment results were visualized using the enrichplot package.

### Biochemistry and Immunocytochemistry

#### Western blotting

Astrocytes were lysed using RIPA buffer (50 mM Tris-HCl, 150 mM NaCl, 0.1% Triton X-100, 0.5% sodium deoxycholate and 0.1% SDS, pH = 7.4) supplemented with Halt Protease and phosphatase inhibitor (ThermoFisher Scientific Cat#78442) at 4 °C. Samples were then clarified via centrifugation at 17,000 × g and supernatant was collected. Total protein concentration per sample was measured using the Pierce BCA Protein Assay Kit (ThermoFisher Scientific, Cat#23225) as per manufacturer’s instructions, and samples were diluted to equal concentrations. Samples were denatured via the addition of denaturing sample buffer (0.4% SDS, 50% glycerol, 125mM tris HCl pH 6.8, 0.2% Orange G) and incubation at 95 °C for 10 minutes.

Samples were resolved via SDS-PAGE. Resolved proteins were transferred to Immobilon-FL PVDF membranes (Millipore) at 100 V. Post transfer, membranes were dried for 60m at RT as per manufacturer’s instructions and re-activated using methanol. For total protein normalization, the membranes were stained with Revert™ 700 Total Protein Stain (Licor Cat#926-11021) for 5 mins, washed once in water and then twice more in water containing 6.7% acetic acid and 30% methanol, and then imaged using the Odyssey CLx Infrared Imaging System (Li-COR). Membranes were then destained with 0.1 M NaOH containing 30% methanol and washed thrice in water and once more in 1x Tris-buffered saline (TBS; 50 mM Tris-HCl, 274 mM NaCl, 9 mM KCl). Membranes were then blocked for 5 mins in EveryBlot Blocking Buffer (BioRad Cat#12010020) and incubated in that same blocking buffer with the indicated primary antibody at 4 °C overnight. The following day, membranes were washed three times with 1× TBS containing 0.1% Tween-20 (TBS-T). Membranes were incubated with secondary antibody (LiCOR) diluted in EveryBlot Blocking Buffer with 0.2% of 10% sodium dodecyl sulfate (10% SDS; BioRad) for 60m at RT, washed thrice more in TBS-T, twice in ddH_2_O, and then imaged in Odyssey CLx Infrared Imaging System. Band intensities were quantified in Image Studio™ Software (Li-COR; Version 5.2.5), normalized to respective total protein intensities, and statistical analyses were performed in Graphpad Prism.

#### Two-step real time quantitative Polymerase Chain Reaction (RT-qPCR)

To quantify transcript levels, 2.75x10^6^ astrocytes per condition were harvested using TRIzol (ThermoFisher, Cat# 15596026) as per the manufacturer’s instructions. cDNA was generated using the SuperScript First-Strand Synthesis System (ThermoFisher, Cat# 11904018) and purified using Zymo Research Oligo Clean & Concentrator kit (Cat# D4060). Each reaction contained 10 ng samples or equivalent volume of nuclease free water (for non-transcript control) and 300 nM of each primer. PowerUP SYBR Green Master Mix (Applied Biosystems, Cat# A25742) was used as instructed to catalyze PCR in a QuantStudio 3 Real-Time PCR System Machine (Applied Biosystems, Cat# A28567), and amplification data was produced with QuantStudio Design and Analysis software.

For each transcript of interest, relative changes in expression were calculated by using a standard 2^-ΔΔCt^ formula and normalizing to *BACT* transcript levels for each sample and then to the negative control in each experiment. Statistical analyses were performed on the resulting 2^-ΔΔCt^ values using Graphpad Prism.

#### Cytokine Assay

To collect samples for the cytokine assay, astrocytes were incubated with 10 μM AA/ 5 μM OA or a vehicle control for eight hours. Cells were then washed twice briefly with hippocampal maintenance media and incubated in hippocampal maintenance media for 16 hours. The amount of maintenance media added to cells for the 16h incubation was kept consistent between conditions (6.5mL for astrocytes plated at a density of 2.75x10^6^). Following this incubation, media was removed from each dish and centrifuged at 3000 rcf to remove any cellular debris. Supernatant was snap frozen in liquid nitrogen and stored at -80 until being sent for analysis via Luminex assay by Eve Technologies. Results of the cytokine assay were imported into R, and outliers were removed if the value fell more than 1.5x outside of the interquartile range for all biological replicates for the condition in question.

#### Immunocytochemistry

For all monoculture immunocytochemistry experiments except the proximity ligation assay, astrocytes (175k) were plated on 35 mm plates with a 20mm glass bottom (MatTek). In experiments intended to visualize Parkin localization to damaged mitochondria, astrocytes were transfected with 0.75 μg human YFP-Parkin using lipofectamine 2000 transfection reagent 24-48h prior to experimentation and fixation. For the representative image shown in Fig. 1K, a TOMM20-halo construct was transfected 24-48 hours before experimentation and fixation, and 100 nM TMR-direct halo ligand (Janelia) was added to cells 2 hours prior to fixation concurrently with the experimental treatment.

Cells were fixed with 4% Paraformaldehyde in PBS (ThermoFisher?) containing 4% sucrose (w/v) for 10 minutes. Samples were then washed thrice with PBS, permeabilized for 5 minutes in 0.25% triton-X in PBS, and blocked for 1 hour in blocking buffer (5% Normal Goat Serum [Sigma Aldrich], 1% Bovine Serum Albumin Fraction V [Fisher Scientific] in PBS, filtered). Primary antibodies were diluted in blocking buffer as indicated in Table S2 and incubated overnight at 4C. Cells were then washed thrice more with 1x PBS and incubated for one hour in AlexaFluor secondary antibody derived from goat (1:500 in blocking buffer) at room temperature. Cells were washed 3 times more in PBS and imaged in PBS.

For proximity ligation assays, 40,000 astrocytes were plated on 7mm glass-bottomed dishes (MatTek). Duolink In Situ PLA Mouse/Rabbit kit with red detection reagents (Sigma Aldrich) was used as directed by the manufacturer, with the addendum that AlexaFluor secondary antibody was added to the amplification step to visualize GFAP. Cells were mounted in the mounting media provided alongside the PLA assay reagents for imaging.

#### Antibodies

Primary antibodies were used at the dilutions as follows: p62/Sqstm1: ICC 1:200, PLA 1:200, immunoblotting 1:750; S65 phosphorylated ubiquitin: ICC 1:250, PLA 1:200, immunoblotting 1:2000; Hsp60: ICC 1:200; GFAP: ICC 1:300; Mic60: ICC 1:100, immunoblotting 1:500; STAT3: immunoblotting 1:750; Phosphorylated (Tyr705) STAT3: immunoblotting 1:500; Phosphorylated JNK/SAPK: immunoblotting 1:500; p38/MAPK: immunoblotting 1:750; Phosphorylated p38/MAPK: immunoblotting 1:500; MAPKAPK2: immunoblotting 1:750; Phosphorylated MAPKAPK2: immunoblotting 1:500; Phosphorylated CREB: immunoblotting 1:500; Mitogen Stress Kinase 1 (MSK1): immunoblotting 1:750; Phosphorylated (Thr581) Mitogen Stress Kinase 1 (MSK1): immunoblotting 1:500; Phosphorylated p65: immunoblotting 1:500; p65: immunoblotting 1:750; LC3: immunoblotting 1:750; Mitofusin 2: immunoblotting 1:1000; Parkin: immunoblotting 1:50; OPTN: immunoblotting 1:2000; TAX1BP1: immunoblotting 1:2000. For information regarding the catalog number and RRIDs of each antibody, please see the key resources table.

### Microscopy and Quantitative Micrograph Analysis

#### Microscopy

Imaging was performed on one of two microscopes. The first was a PerkinElmer UltraView Vox spinning disk confocal on a Nikon Eclipse Ti Microscope using an apochromatic 100x 1.49 NA oil immersion objective. Images on this microscope were captured using a Hamamatsu CMOS ORCA-Fusion (C11440-20UP). The second microscope was a Leica DMi6000 stand outfitted with a Crest Optics X-Light V3 confocal unit. Images on this microscope were acquired using an apochromatic 100x 1.49 NA oil immersion objective and a pco.edge 4.2 bi back-illuminated sCMOS camera.

Z-stacks were collected at 200nm increments through the full volume of the cell. Both microscopes utilize VisiView (Visitron) operating software. All biological replicates for each experiment were imaged on the same microscope, and illumination settings were kept consistent for each replicate to minimize variability.

#### Image analysis

All image analysis aside from the neuronal mitochondrial volume measurements was performed in Fiji Is Just ImageJ (FIJI; version 2.14.0/1.54f). To analyze mitochondrial intensity of YFP-Parkin and phospho-ubiquitin in mono-cultured and co-cultured astrocytes and TMRE intensity in hippocampal neurons, mitochondrial ROIs were generated based on the mitochondrial marker (Mic60 for astrocytes, mito-mEmerald for neurons) using a rolling ball background subtraction (radius 30) followed by an automated otsu thresholding step and a processing step applying the despeckle function to avoid any inclusion of noise. ROIs were then overlaid on the channel of interest (YFP-Parkin, phospho-ubiquitin, TMRE) and mean intensity within the mitochondrial bounds was quantified. Mitochondrial fragmentation was quantified via the same segmentation method and calculation of average mitochondrial area per cell. Quantifications were done on maximum projections of the relevant images. To quantify interactions between phospho-ubiquitin and p62 using a proximity ligation assay (PLA), images of the ligation assay indicating p62/phospho-ubiquitin interactions were segmented using Ilastik machine learning software. Cells were manually traced based on GFAP fluorescence and puncta number per cell was measured using the analyze particles feature for each cell in FIJI.

To analyze p62 puncta per cell, astrocytes were manually traced to define cellular area, and the p62 channel was segmented using Ilastik machine learning software. Puncta area was measured using the analyze particles feature in FIJI and the number of puncta in each cell was normalized to cell area. To analyze the percent of p62 that localized to phospho-ubiquitin, the respective phospho-ubiquitin images were also segmented in Ilastik and the Boolean “and” function in the FIJI image calculator was applied to p62 and phospho-ubiquitin segmentations from the same channel to identify areas of p62/phospho-ubiquitin overlap. The area of overlap was then divided by the total area for p62 puncta for each cell and multiplied by 100 to generate a percentage for p62 localized to phospho-ubiquitin. Quantifications were performed using maximum projections.

Mitochondrial volume analysis in hippocampal rat neurons was performed in Imaris. The neuronal soma was manually segmented based on fluorescence in the Beta-III-tubulin channel using manual isometric contouring. Mitochondria were segmented into surfaces using automated thresholding with background subtraction, with the diameter of the largest sphere that fits into the object set as 0.487 μm. Non-somal mitochondria were filtered out in a manner contingent on overlap with the somal surface; mitochondrial objects whose volume overlapped with the soma at a ratio of greater than 0.4 were analyzed. Due to variability in three-dimensional analysis, all volumetric data for individual somas were pooled and outliers (5 total, 3 from the control and 2 from the TNF-α-treated condition) were removed via application of a standard Robust Regression and Outlier Removal (Q=1%) in Graphpad Prism. Statistics were run on the average mitochondrial volumes per soma for the independent biological replicates. All statistical analyses on imaging data were performed in Graphpad Prism.

#### Experimental data availability

Both the raw data files associated with this study and the source data will be uploaded to Zenodo prior to or upon formal acceptance for publication.

## KEY RESOURCES TABLE

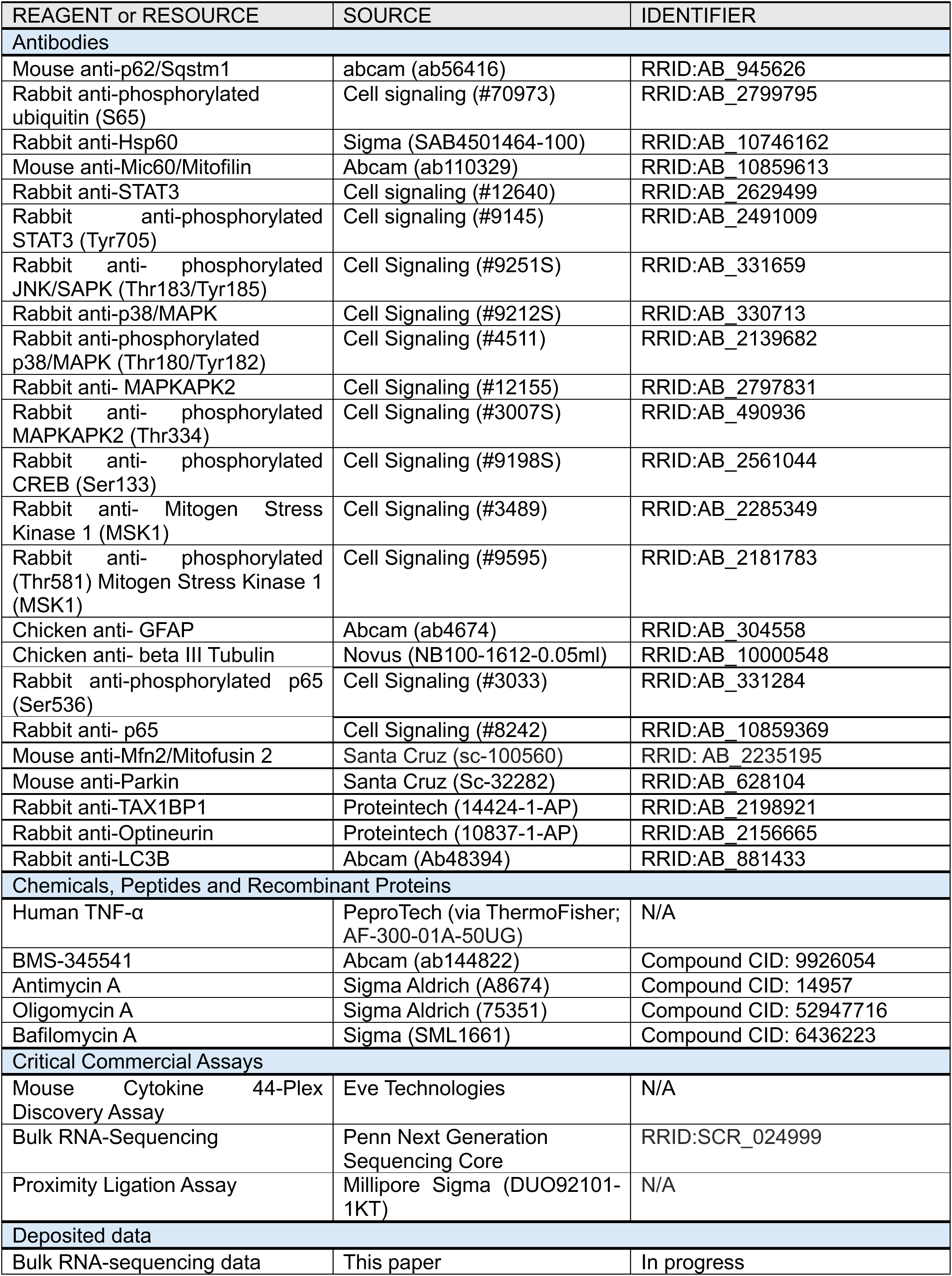
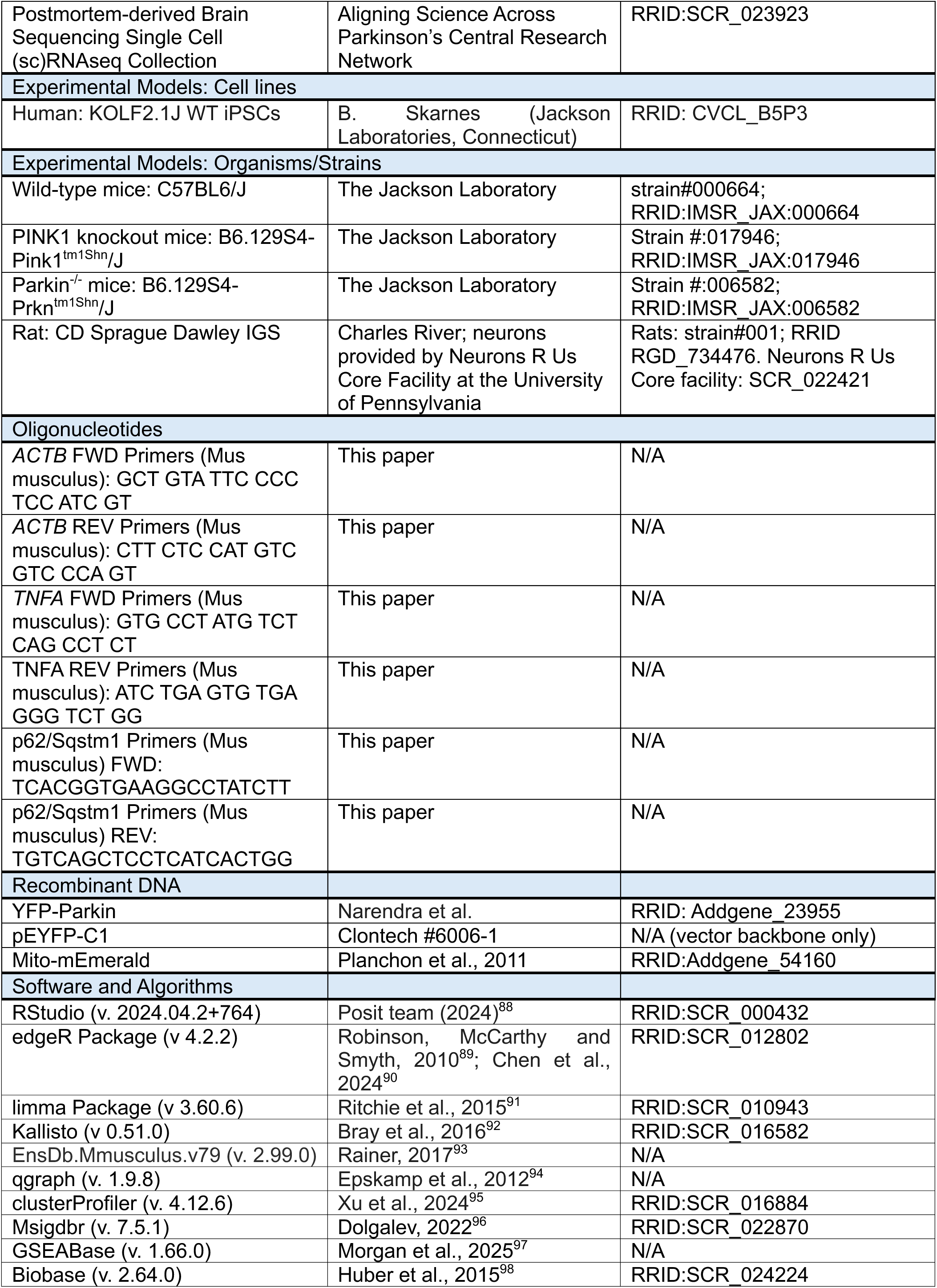
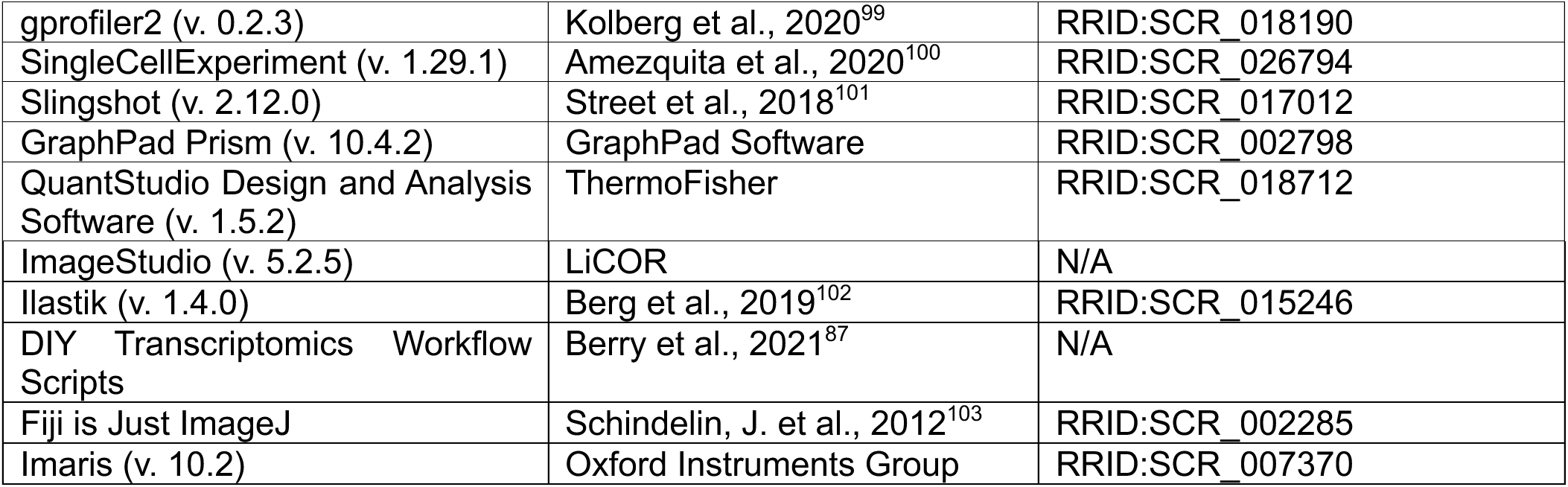

---

**Extended Data Figure 1.**
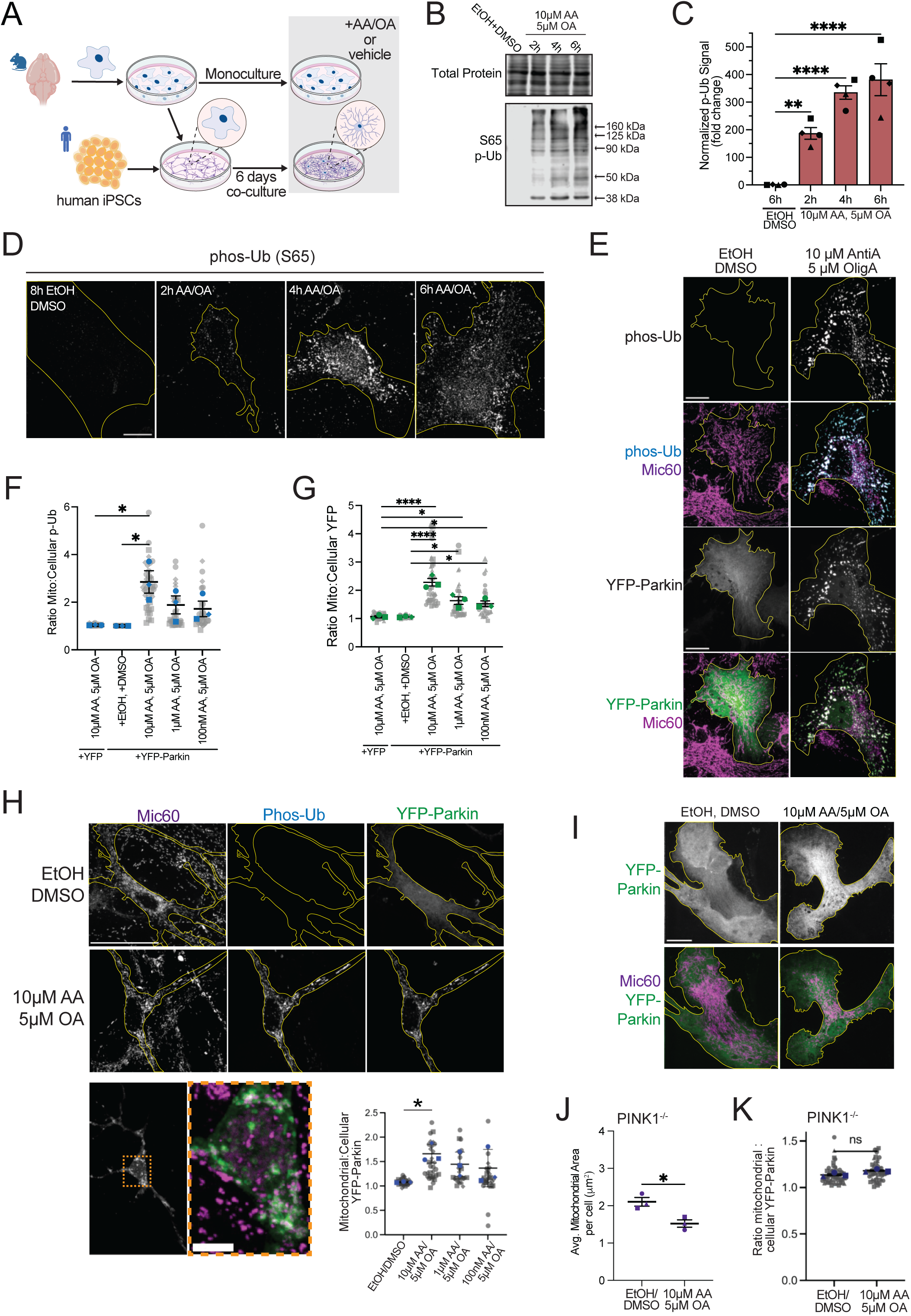
OXPHOS inhibitors induce mitochondrial fragmentation and PINK1/Parkin-dependent mitophagy in astrocytes. A) Schematic illustrating primary astrocyte monoculture and co-culture with human iPSC-derived neurons. B, C) Western blots detecting S65 phospho-ubiquitin in monoculture astrocytes following 10 μM AA/ 5 μM OA treatment for 2h, 4h, and 6h, or 6h of vehicle (EtOH/DMSO) control, and corresponding quantitation (n=4). Data analyzed by one-way ANOVA, error bars are mean +/- SEM. D) ICC for phospho-ubiquitin (S65) following 2, 4, or 6 hours of treatment with 10 μM AA/ 5 μM OA or a vehicle control. Scale bar = 20 μm. E-G) Immunostaining in monoculture astrocytes to detect S65 phospho-ubiquitin and mitochondrial protein Mic60 (E, F) or YFP-Parkin recruitment to damaged mitochondria using the same marker (E, G) following 2h of treatment with various doses of AA/OA. Representative max projections shown. Gray dots represent individual cell measurements; blue dots represent average for each biological replicate (n=3, ordinary one-way ANOVA, error bars are mean +/- SEM). Scale bar =20 μm. H) Astrocytes transfected with YFP-Parkin at 6 days post-co-culture with iNeurons, treated with 2h 10 μM AA/ 5 μM OA or a vehicle control and immunostained for Mic60 and phospho-ubiquitin (S65). Ratio of mitochondrial YFP-Parkin intensity: total cellular intensity was quantified (n= 3, ordinary one-way ANOVA with multiple comparisons, error bars are +/- SEM). Scale bar = 20 μm; inset scale bar= 5 μm. I-K) Representative images (I) of average mitochondrial area per cell (J) and ratio mitochondrial YFP-Parkin intensity: cellular YFP-Parkin intensity (K) in PINK1^-/-^ astrocytes treated with 2 hours 10 μM AA/ 5 μM OA or a vehicle control. Representative images are max projections (n=3, unpaired t-test, error bars are +/- SEM). Biological replicates are defined as independent experiments performed on astrocytes from different dissections. For all graphs, p ≤ 0.05 = *; p < 0.01 = **; p < 0.001 = ***; p < 0.0001 = ****.

**Extended Data Figure 2.**
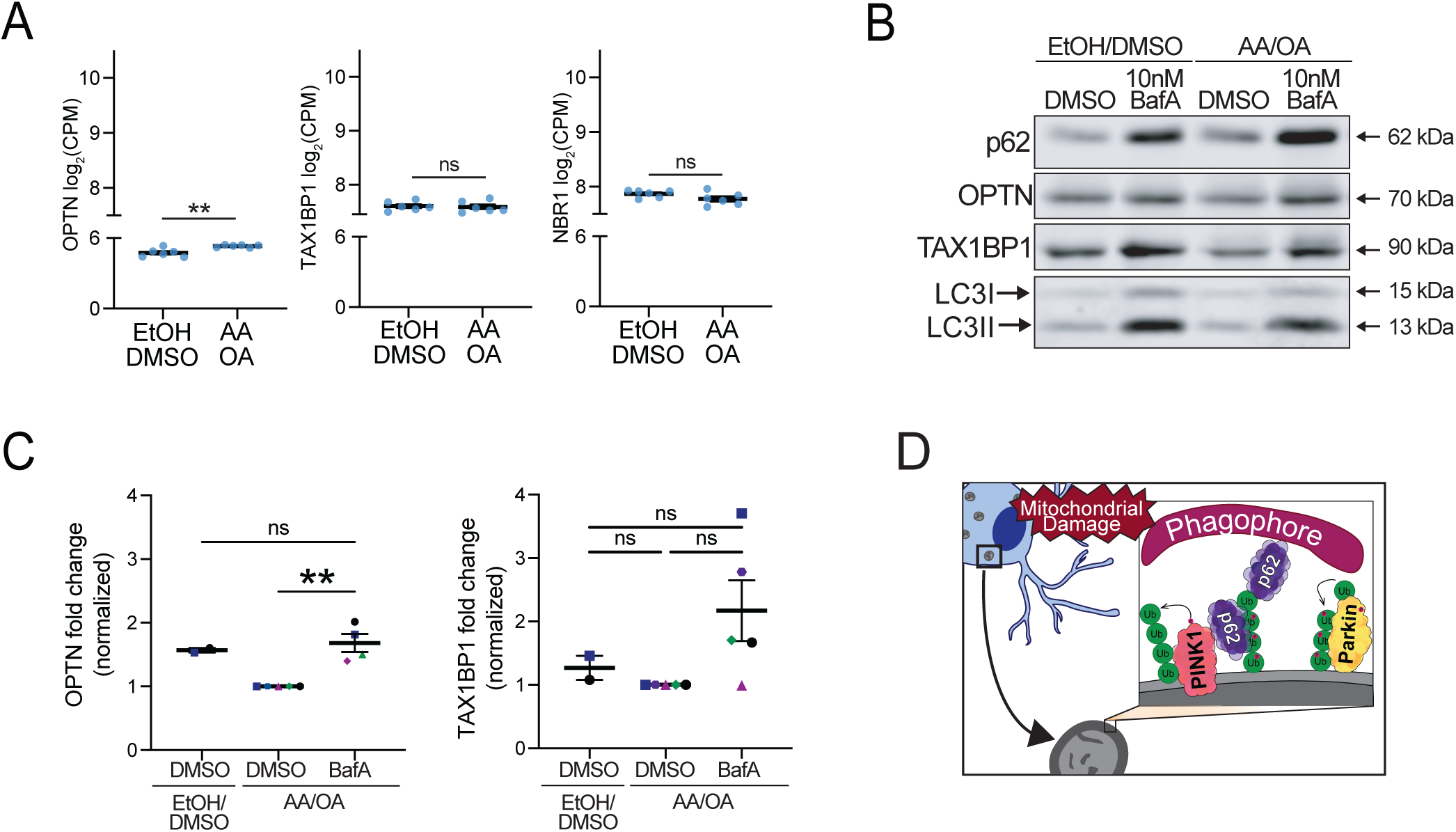
Ubiquitin-binding autophagy receptors OPTN, TAX1BP1, and NBR1 show minimal expression changes in response to mitochondrial damage or mitophagic flux assays. A) Log_2_CPM for OPTN, NBR1, and TAX1BP1- the non-p62 autophagy receptors that contain ubiquitin binding domains and were detectable via bulk RNA-seq- in WT astrocytes post-stress compared to the control (n=6; unpaired t-tests; error bars indicate mean +/- SEM). B) Representative western blots for autophagy receptors in astrocytes treated for 16 hours with either 10 nM Bafilomycin A (BafA) or a vehicle control in conjunction with either 10 μM AA/ 5 μM OA or a vehicle control. C) Quantifications for the western blots shown in E for OPTN and TAX1BP1. Each replicate is normalized to the AA/OA-only treatment condition (n=2 for treatment with both vehicle controls, n=5 for DMSO vs. BafA comparison following AA/OA; ordinary one-way ANOVA with multiple comparisons, error bars are mean +/- SEM). D) Schematic of PINK1/Parkin mitophagy and p62 recruitment.

**Extended Data Figure 3.**
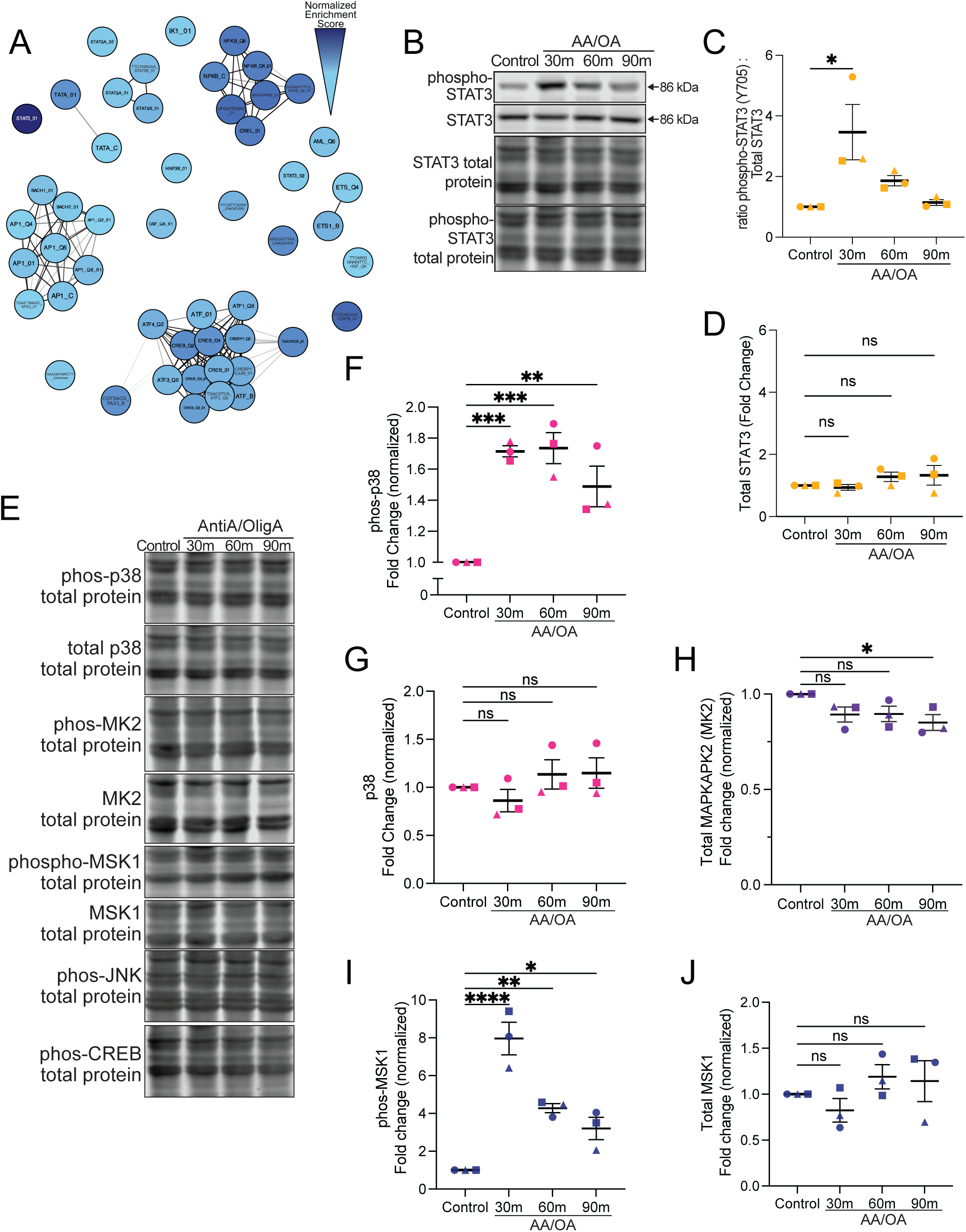
MSK1 and p38 are phosphorylated, but not upregulated, following mitochondrial damage to astrocytes. A) GSEA results for the 50 most highly enriched TFT-legacy terms in WT astrocytes following 8h 10 μM AA/ 5 μM OA treatment compared to vehicle control. Terms with higher overlap in their defining genes are proximal to each other. Color is based on normalized enrichment score relative to other terms in the graph. Results based on average gene expression changes across 6 independent biological replicates. B) Representative western blots for STAT3 and phospho-STAT3 in astrocytes following 30, 60, or 90 minutes of 10 μM AA/ 5 μM OA treatment or 90 minutes of treatment with a vehicle control. C, D) Western blot quantifications of (C) the ratio of phospho-STAT3 (Y705) : total STAT3 and of (D) total STAT3 following 30, 60, or 90 minutes of 10 μM AA/ 5 μM OA treatment or 90 minutes of treatment with vehicle control (n= 3, ordinary one-way ANOVA with multiple comparisons, error bars are mean +/-SEM). E) Total protein stains for each of the representative blots in Fig. 4D. F, G) Raw levels of phosphorylated p38 (F) or total p38 levels (G) normalized to respective biological replicate vehicle controls following 30m, 60m or 90m of 10 μM AA/ 5 μM OA treatment or 90m treatment with a vehicle control (n=3, ordinary one-way ANOVA, error bars are mean +/- SEM). H) Total levels of MAPKAPK2 (normalized to respective biological control) following 30m, 60m or 90m of 10 μM AA/ 5 μM OA treatment or 90m treatment with a vehicle control (n=3, ordinary one-way ANOVA, error bars are mean +/- SEM). I) Levels of phospho-MSK1(T581; normalized to respective biological control) following 30m, 60m or 90m of 10 μM AA/ 5 μM OA treatment or 90m treatment with a vehicle control (n=3, ordinary one-way ANOVA, error bars are mean +/- SEM). J) Total levels of MSK1 (normalized to respective biological control) following 30m, 60m or 90m of 10 μM AA/ 5 μM OA treatment or 90m treatment with a vehicle control (n=3, ordinary one-way ANOVA, error bars are mean +/- SEM).

**Extended Data Figure 4.**
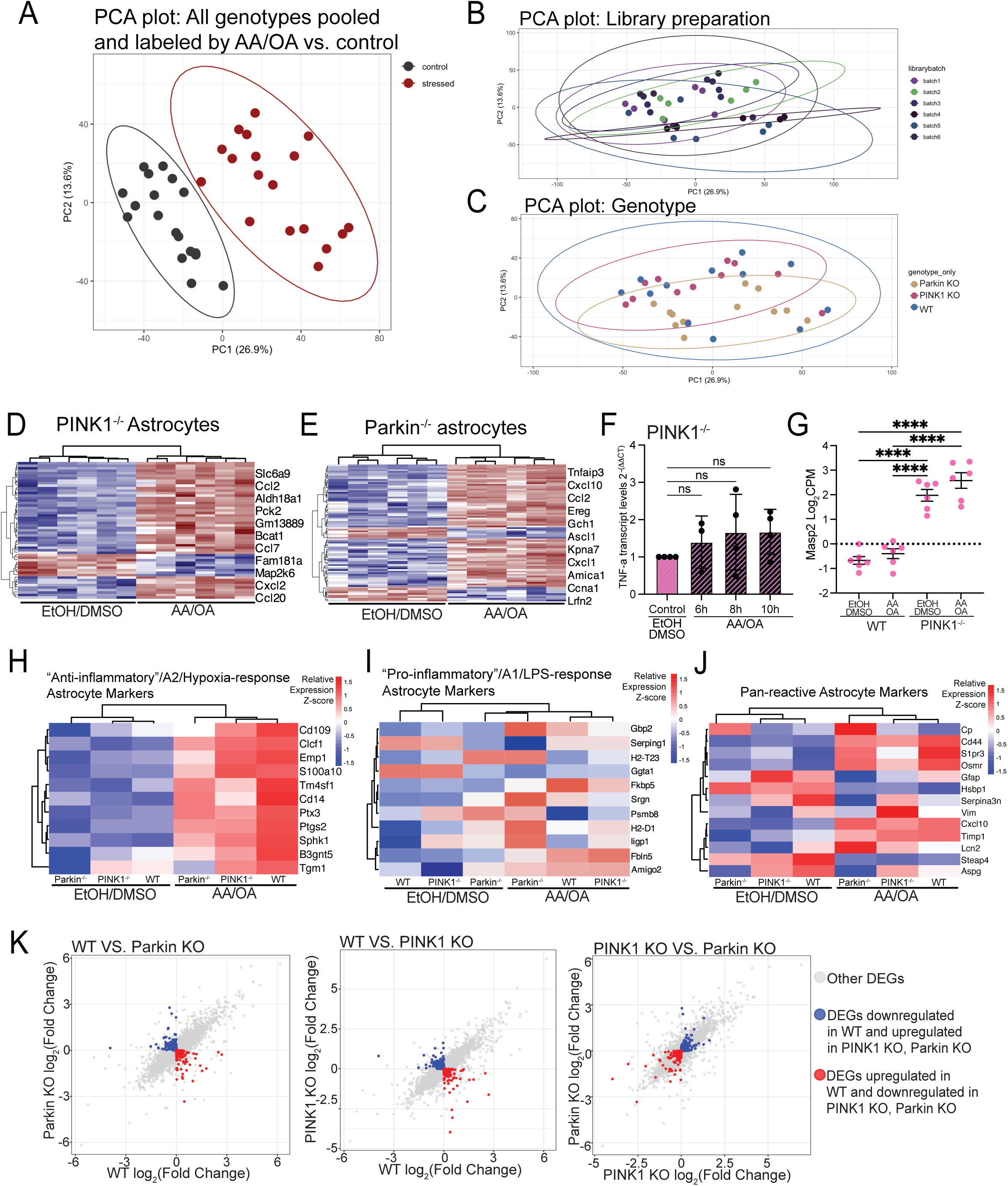
PINK1-/-, Parkin-/-, and WT astrocytes have different DEGs following OXPHOS inhibitor treatment. A) Principal component analysis pooling all biological replicates of astrocytes following 8h 10 μM AA/ 5 μM OA treatment (denoted as stressed) and all biological replicates treated with a vehicle control. Ellipses indicate 95% confidence regions based on a multivariate t-distribution (AA/OA, n=18; control, n=18). B) Principal component analysis pooling all biological replicates in the bulk RNA-sequencing experiment by the date of library preparation. Ellipses indicate 95% confidence regions based on a multivariate t-distribution. C) Principal component analysis pooling all biological replicates in the bulk RNA-sequencing experiment by the genotype of the astrocytes. Ellipses indicate 95% confidence regions based on a multivariate t-distribution. D,E) Differentially expressed genes (DEGs) between PINK1^-/-^ astrocytes (D) and Parkin^-/-^ astrocytes (E) following 8h treatment with 10 μM AA/ 5 μM OA or vehicle control. Columns represent biological replicates (n=6 for each). DEGs were identified using limma with a Benjamini–Hochberg correction (adjusted p < 0.01, |log2FC| ≥ 2). Expression values scaled by row. Genes and samples were hierarchically clustered using correlation-based distance metrics. F) qPCR measuring transcript levels for *Tnfa* normalized to *Actb* following 6h, 8h, or 10h treatment with 10 μM AA/ 5 μM OA or 10h treatment with vehicle in PINK1^-/-^ astrocytes (n= 4, non-parametric ANOVA, error bars indicate mean +/- SEM). G) Transcript levels of mannan-binding lectin serine protease 2 (MASP2) in WT and PINK1^-/-^ astrocytes treated with 10 μM AA/ 5 μM OA or a vehicle control for 8h (n=6, ordinary one-way ANOVA, error bars are mean +/- SEM). H, I, J) Heatmaps comparing A2 astrocyte markers (G), Pan-reactive astrocyte markers (H), and pro-inflammatory astrocyte markers (I) across WT, PINK1^-/-^, and Parkin^-/-^ astrocytes following either 8h 10 μM AA/ 5 μM OA treatment or treatment with a vehicle control. Columns represent average expression across biological replicates (n=6 per condition). Expression values are scaled by row to visualize relative expression. K) DEGs in WT, PINK1^-/-^ and Parkin^-/-^ astrocytes following 8h treatment 10 μM AA/ 5 μM OA compared directly to one another. In each scatter plot, the DEGs that were downregulated in both knockouts and upregulated in WT (red) or downregulated in WT and upregulated in both knockouts (blue) are indicated. Biological replicates are defined as independent experiments performed on astrocytes from different dissections. For all graphs, p < 0.05 = *; p < 0.01 = **; p < 0.001 = ***; p < 0.0001 = ****.

**Extended Data Figure 5.**
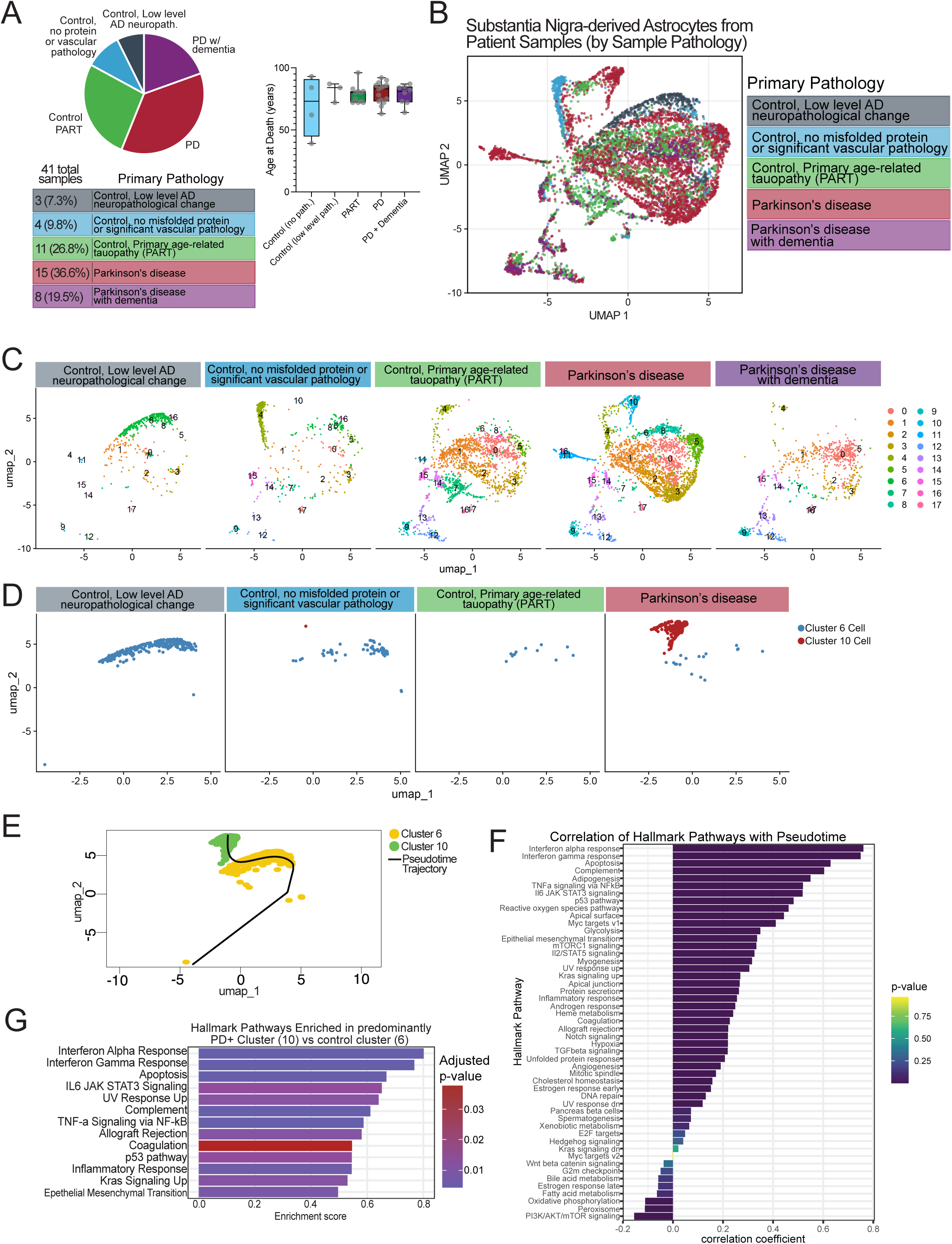
scRNA sequencing analysis revealed clusters enriched for control- and PD-derived astrocytes. A) Overview of tissue samples used in the experiments that generated the scRNA-sequencing data utilized for analysis of cells annotated as astrocytes from the substantia nigra of human brain tissue. Pie chart and table (left) indicates the number of samples corresponding to each primary pathology; box and whiskers plot (right) represents the age at death for each patient from whom the sample was collected (box represents median and interquartile range). B) Seurat-generated cluster analysis in Fig. 6A colorized by primary pathology of the sample that each cell was derived from. C) UMAP plots of the cells from each primary pathology organized into the seurat-generated clusters shown in Fig. 6A to illustrate distribution of cells from different pathologies across clusters. D) UMAP plots including only cells that were sorted into the two Seurat-generated clusters of interest. Each plot includes cells in these clusters that originated from tissue annotated as having the primary phenotype designated. The “Parkinson’s disease with dementia” primary pathology was excluded because there were no astrocytes derived from samples categorized as being from samples with that pathology that were in either of these clusters. E) The inferred pseudotime trajectory curve from control-enriched cluster to the PD-enriched cluster, pictured on the original UMAP plot in which the clusters were defined. F) Bar plot showing Pearson correlation coefficients (r) between Hallmark pathway module scores and inferred pseudotime. P-values (indicated by bar color) were obtained from two-sided Pearson correlation tests, with the null hypothesis that the correlation between module score and pseudotime equals zero. G) Bar graph summarizing Hallmark Gene Set Enrichment Analysis results for the comparison between astrocyte clusters 10 and 6. Bars represent enrichment scores derived from GSEA, with higher positive values indicating stronger enrichment in cluster 10. Bar color denotes the adjusted *p*-value for each gene set.

**Extended Data Figure 6.**
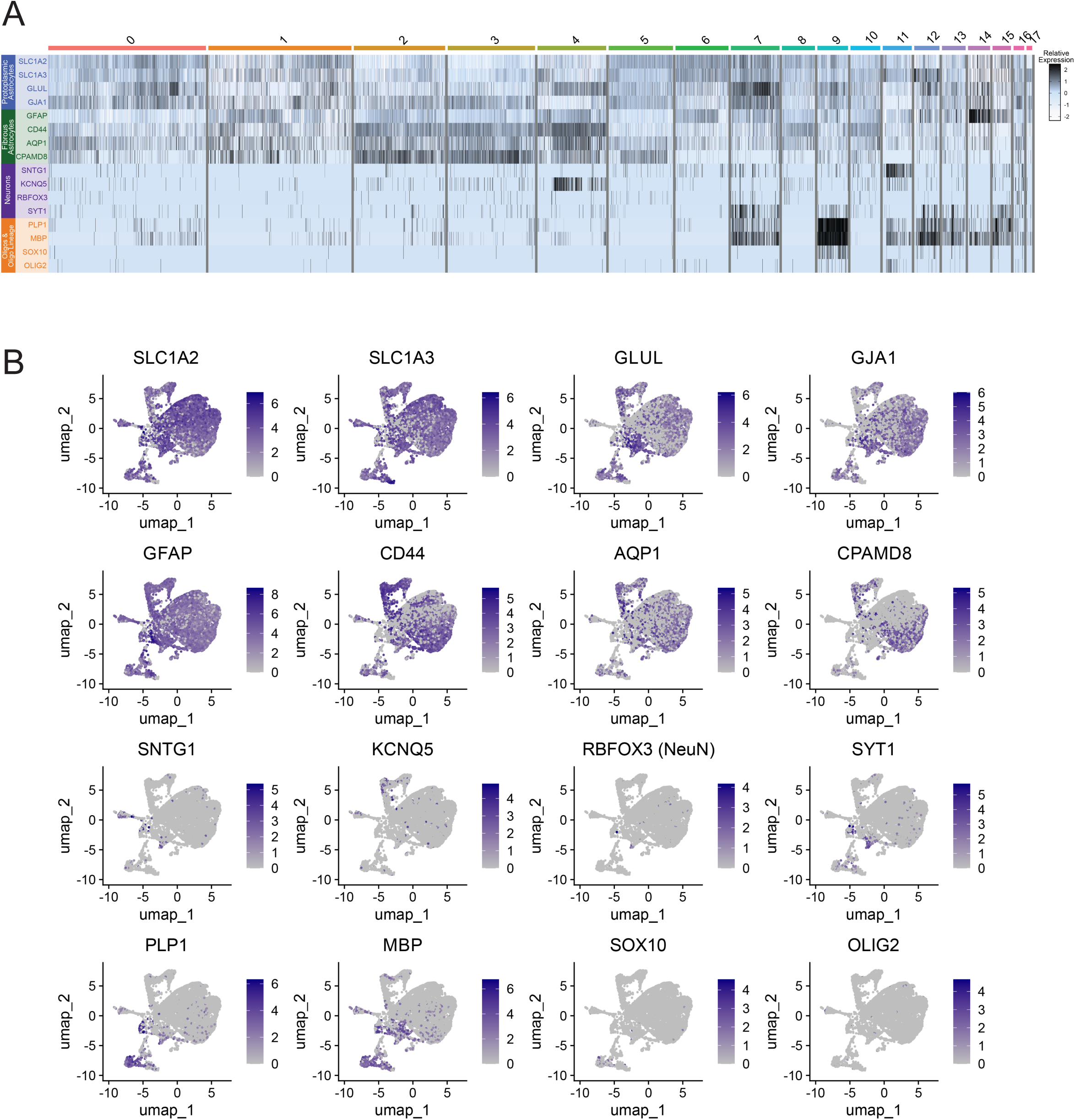
Validating astrocytic character of cells used for scRNA-sequencing analysis. A) Heatmap showing relative expression of genetic markers for protoplasmic astrocytes (*SLC1A2, SLC1A3, GLUL, GJA1*), fibrous astrocytes (*GFAP, CD44, AQP1, CPAMD8*), neurons (*SNTG1, KCNQ5, RBFOX3, SYT1*) and oligodendrocyte lineage cells (*PLP1, MBP, SOX10, OLIG2*) across cells annotated as astrocytes, embedded in UMAP space. Colors represent scaled gene expression values, where expression for each gene was centered and scaled across all cells (z-scoring). Cells are grouped by Seurat cluster identity, genes are displayed as rows, and individual cells are shown as columns. B) Feature plot depicting relative expression of the same 16 markers of interest for the four different cell types described in (A). Color scales represent log-normalized gene expression values.

**Supplementary Table 1.**
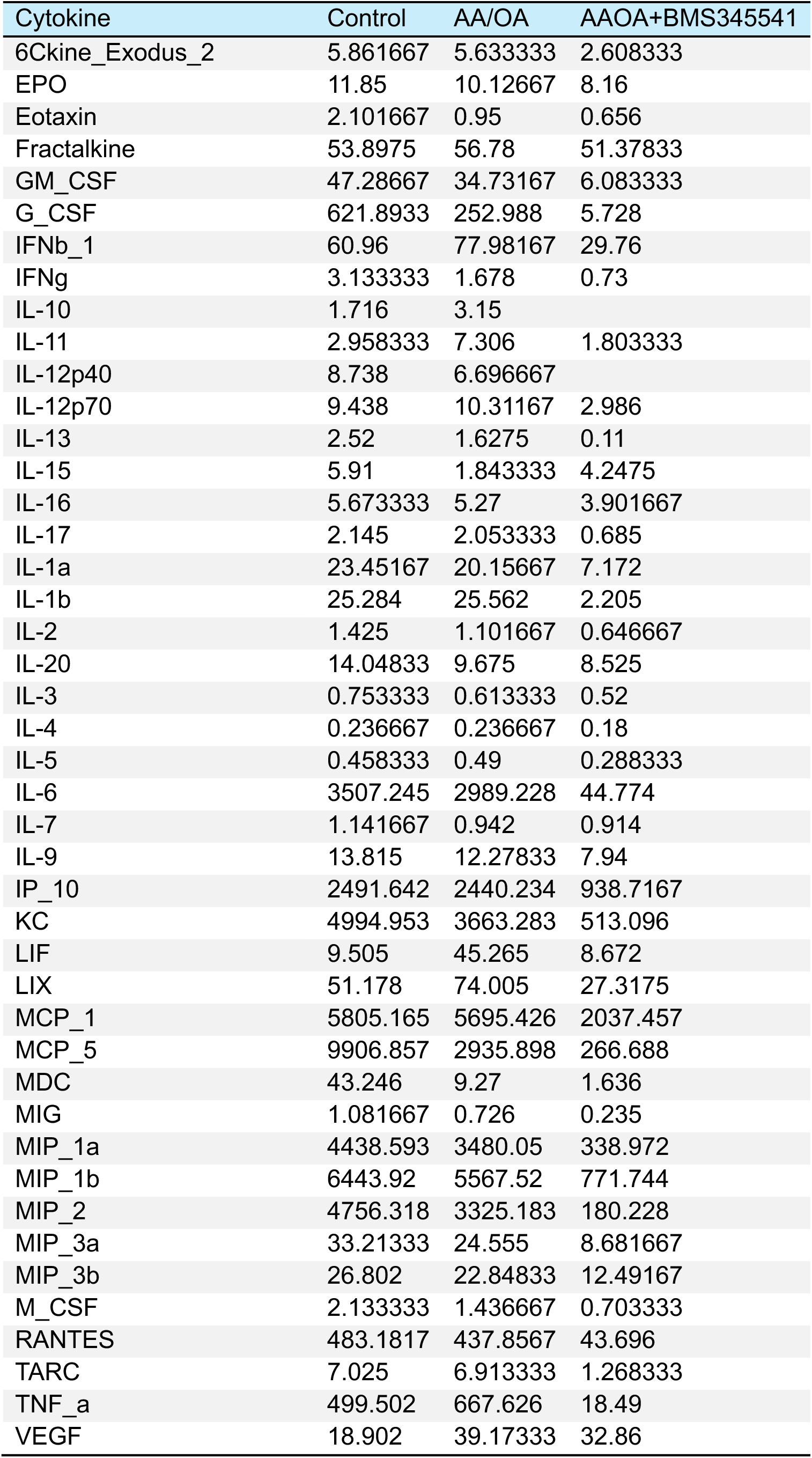
Raw concentrations of secreted factors (pg/mL) detected in media following treatment with 5 μM BMS345541 or vehicle control for 72h, as well as OXPHOS inhibitors (10 μM Antimycin A, 5 μM Oligomycin A) or vehicle control for the final 16h of treatment.

